# Intestinal molecular siphoning by engineered bacterial spores for non-systemic treatment of nicotine addiction

**DOI:** 10.64898/2026.11.03.763521

**Authors:** Xisheng Xiong, Ruiqi Zhu, Yueqing Han, Mengxin Yu, Xuhui Xiang, Yulan Shao, Yijun Chen

## Abstract

Eradicating toxic small molecules from the body typically requires the administrations of therapeutic agents for systemic circulation and tissue distribution, resulting in inevitable adverse events from off-target effects. Here, we propose a “intestinal molecular siphoning” approach for treating diseases caused by toxic substances, in which enzymatic degradation of a toxic substance in the intestinal lumen serves as a driving force for the diffusion of the toxic substance from blood to gut, and continuous elimination of the toxic substance in the gut could effectively ease substance dependence. Using nicotine as a model substance, a live bacterial system was constructed by displaying a trypsin-resistant nicotine oxidase on the surface of *Bacillus subtilis* spores, and encapsulating in enteric-coated microspheres for gastric protection and targeted intestinal release. After oral treatments with the live bacterial system in mice, peak concentrations of nicotine in blood and brain were remarkably reduced compared to controls. Moreover, the live bacterial system could effectively attenuate nicotine addiction by suppressing dopamine reward surges to reverse withdrawal hyperalgesia, preventing the upregulation of nicotinic acetylcholine receptors, and alleviating anxiety-like behaviors. These findings show the potential of “intestinal molecular siphoning” as a target-independent therapeutic strategy for lowering circulating toxic substances through non-systemic enzyme reprogramming.

**Significance Statement:** Currently, blood and tissue exposures are a critical measure on the efficacy for conventional therapeutics, including small molecules, biologics or therapeutic cells. To avoid various off-target-associated adverse events from such exposures, bypassing bloodstream to exert systemic therapeutic effects is a prerequisite to achieve target-independent treatments. Here, we describe the feasibility of implementing “intestinal molecular siphoning” approach for degrading toxic molecules in the body. When nicotine oxidase is displayed on the surface of probiotic *B. subtilis*, oral administration of the engineered live bacterial system exhibits desired efficacy and safety for nicotine addiction in murine models. Beyond the treatment of nicotine addiction, this approach also provides a versatile platform for non-systemically reducing other toxic molecules in the circulations.

## Introduction

Many diseases are caused by harmful small molecules circulating in the bloodstreams, including addictive substances such as nicotine, uremic toxins and drug metabolites. Currently, therapeutic strategies for treating diseases generally require systemic administration or oral absorption of therapeutic agents, which usually results in various adverse events primarily due to wide distribution and tissue residence (1, 2). To avoid such undesired effects, the therapeutic agents should be administrated through a different route that can bypass the bloodstream while maintain the efficacy. Unfortunately, existing therapeutic options are unable to fulfill this demand. Therefore, the development of a feasible approach to non-systemically remove circulating toxic molecules would be advantageous, especially regarding safety concerns.

The intestine, possessing a large absorptive surface area and extensive vascularization, provides a bidirectional interface between the gut lumen and systemic circulation. In most cases, small molecules can passively diffuse from the blood into the intestinal lumen and, in principle, be captured and degraded locally. A recent study showed that nicotine is accumulated at high levels in the ileum during smoking (3), suggesting that the intestine may serve as a reservoir for certain circulating molecules. We therefore hypothesized that continuous degradation of a target molecule by an enzymatic system within the gut lumen would create a concentration gradient to drive a faster diffusion from the bloodstream into the intestine. This process would establish a “molecular siphon” that further promotes the enzymatic degradation and removal of systemic toxic molecules through intestinal excretion. By restricting the degradation system in the gut lumen, the catalytic system could achieve a target-independent therapeutic outcome, thereby eliminating most adverse events, including systemic immunogenicity, receptor bindings, off-target effects, and nonspecific tissue exposures.

Tobacco smoking remains a leading preventable cause of death worldwide (2, 4), yet current smoking-cessation therapies have only shown a limited efficacy. Nicotine, the primary addictive component in tobacco, drives physiological and psychological dependence through central reward pathways (1, 5). Varenicline, one of the most effective pharmacotherapies for smoking cessation, acts as a partial agonist of nicotinic acetylcholine receptor (nAChR). However, its use is largely limited by adverse events, particularly neuropsychiatric symptoms (6). Meanwhile, the discovery of a bacterial nicotine-degrading enzyme NicA2 has shed a new light on smoking cessation, in which nicotine in the circulation could be enzymatically degraded by NicA2 (7, 8). Although intravenous administration of NicA2 enzyme at high doses of 10–70 mg/kg could effectively lower nicotine levels in rodent bloodstreams, strong immunogenicity and repeated intravenous administration prevent it as a therapeutic agent (8). Subsequently, different engineering approaches, such as fusion with albumin-binding domains (9, 10) and PEGylation of the protein (11), are implemented and evaluated, aiming to improve the druggability of NicA2. Unfortunately, the inherited issues of a bacterium-derived protein, especially immunogenicity, are still far from druggable.

In this study, we developed a live bacterial system based on intestinal molecular siphoning. Using nicotine as the substrate, we identified and engineered a trypsin-resistant nicotine oxidase (ssNox) and then displayed it on the surface of probiotic *Bacillus subtilis* spores for directly oxidizing nicotine in the intestine and diffused from blood. The engineered spores were subsequently encapsulated as pH-responsive enteric microspheres for targeted release in the intestine. The combination of surface-displayed ssNox and enteric encapsulation could retain the catalytic machinery within the gastrointestinal tract while continuously degrade luminal nicotine with rapid onset and high efficiency. After the evaluations on the aspects of therapeutic efficacy, neurochemical changes, behavioral differences, and toxicities in murine models, the microspheres with the function of intestinal molecular siphoning were demonstrated to reduce systemic nicotine levels and disrupt nicotine-dependence cycle, which was further confirmed by a favorable safety profile in mice. Collectively, this target-independent approach showed the potential as a new therapeutic option.

## Results

### Engineering of a trypsin-resistant nicotine oxidase for intestinal degradation

Since the catalytic performance of nicotine oxidase determines the outcomes of intestinal degradation and the strength of molecular siphoning, an enzyme capable of retaining activity in the protease-rich intestinal environment is essential. To achieve this, we first used NicA2 as a query for homology searching to obtain a previously uncharacterized NicA2 homologous protein from *Stutzerimonas stutzeri*, which was designated ssNox (Fig. 1A). After cloning, heterologous expression and purification, we found that wild-type (WT) ssNox was rapidly inactivated by low concentrations of trypsin (Fig. S1), indicating that protease resistance is required for intestinal application.

**Figure 1.**
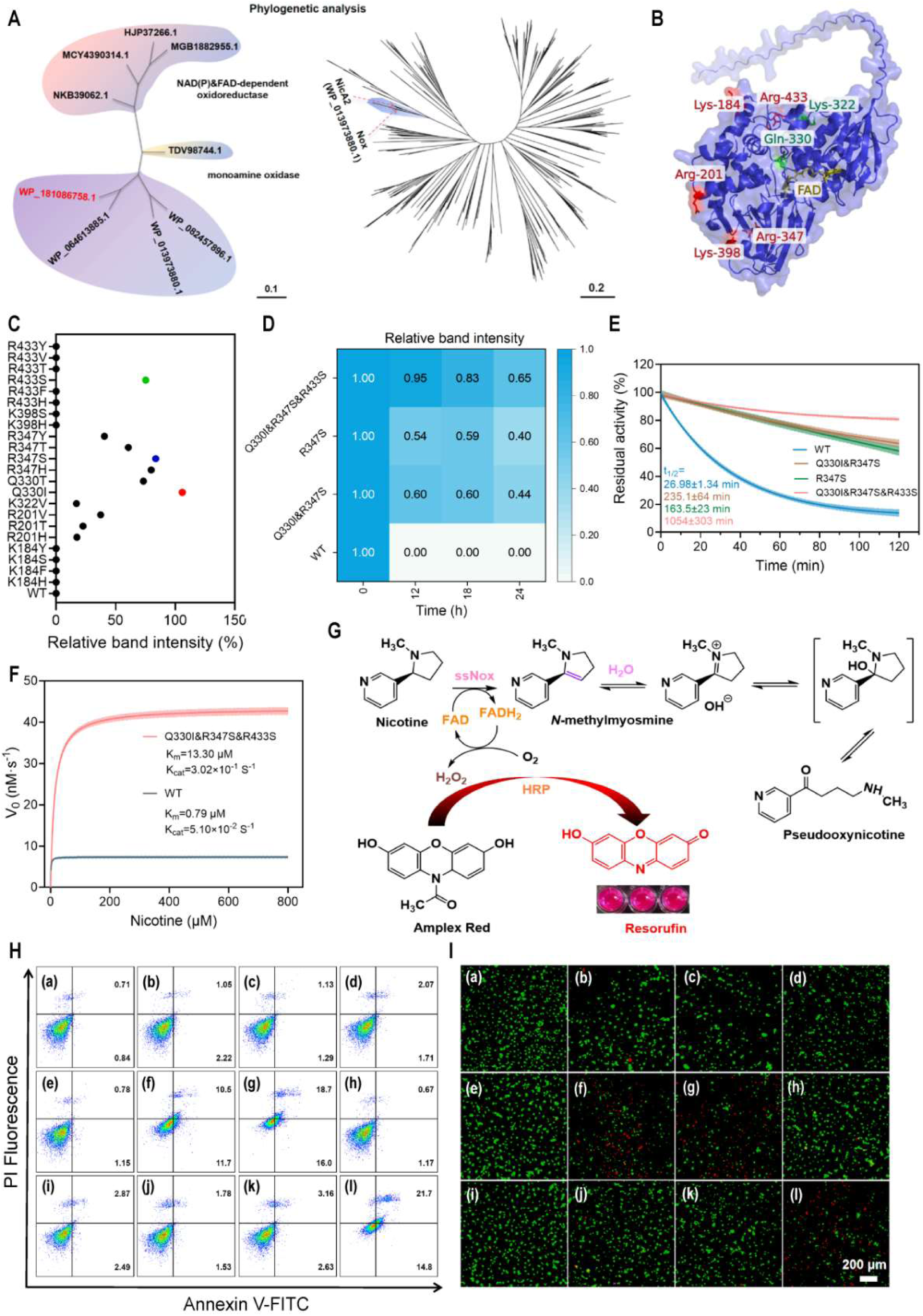
Engineering of a trypsin-resistant nicotine oxidase for intestinal catalysis. (A) Phylogenetic analysis of bacterial homologs of NicA2 with ssNox highlighted. (B) Structural model of ssNox showing the sites selected for engineering. (C) SDS-PAGE analysis of the integrity of mutant proteins after the treatment with trypsin for 4 h at a protein-to-trypsin mass ratio of 1:200. (D) Protein integrity of iteratively optimized mutants after extended trypsin treatment for 12 to 24 h. (E) Residual activity of the triple mutant Q330I&R347S&R433S during trypsin treatment. (F) Kinetic parameter comparison of the mutant Q330I&R347S&R433S and WT ssNox (n=3). (G) Schematic representation of ssNox-catalyzed nicotine oxidation and H_2_O_2_ detection. (H) Flow cytometric analysis of apoptosis in IEC-6 cells following co-incubation with ssNox and nicotine. (I) Calcein-AM/PI viability assay showing H_2_O_2_-associated cytotoxicity and its attenuation by catalase. Panels (a)–(l) represent the following treatments: (a) PBS; (b) 3 ng/mL nicotine; (c) 30 ng/mL nicotine; (d) 300 ng/mL nicotine; (e) 3 ng/mL nicotine and 0.1 mg/mL ssNox; (f) 30 ng/mL nicotine and 0.1 mg/mL ssNox; (g) 300 ng/mL nicotine and 0.1 mg/mL ssNox; (h) 300 ng/mL nicotine and 0.1 mg/mL ssNox in the presence of 5 μg/mL catalase, (i) 300 ng/mL nicotine and 5 μg/mL catalase; (j) 0.1 mg/mL ssNox alone; (k) 5 μg/mL catalase alone, (l) 100 μM H_2_O_2_.

Next, based on structural predictions by AlphaFold3 (12) and known cleavage preferences of intestinal trypsin (13, 14), we performed semi-rational mutagenesis at possible trypsin cleavage sites (Fig. 1B). Iterative mutagenesis followed by the increase of trypsin challenge for 4, 12, 18 and 24 h yielded the best mutant Q330I&R347S&R433S. More than 60% of the mutant protein remained full-length after 24 h of trypsin treatment (Fig. 1C, 1D), corresponding to a half-life of 1,054 min, which is approximately 39-fold longer than of WT ssNox (27 min) (Fig. 1E). The mutant exhibited a higher catalytic turnover number, with a *k_cat_* of 0.302 s⁻¹. Compared to 0.051 s⁻¹ by WT ssNox and previously reported value of 0.013 s⁻¹ by NicA2 (7), the catalytic efficiency was drastically increased (Fig. 1F). Regarding substrate inhibition, the reaction rate by WT ssNox reached a plateau at nicotine concentrations below 1 μM, whereas the mutant maintained nicotine-degrading activity at higher substrate concentrations.

Because nicotine oxidation by ssNox generates H_2_O_2_ as a by-product (Fig. 1G), we assessed the effects of enzymatically generated H_2_O_2_ on IEC-6 cell line derived from intestinal epithelial cells. Flow cytometry and live/dead staining showed that the amount of H_2_O_2_ produced during nicotine degradation increased the levels of apoptosis and necroptosis comparable to those induced by a direct treatment with 100 μM H_2_O_2_ (Fig. 1H, 1I; Fig. S4). By contrast, nicotine alone at physiologically relevant concentrations of 3–300 ng/mL did not induce apoptosis or necrosis in IEC-6 cells. Catalase supplementation indeed abolished these toxic effects, suggesting that H_2_O_2_ detoxification would be necessary for intestinal application of ssNox. Together, a desired mutant of bacterial nicotine oxidase was generated, and its intestinal use would require the decomposition of by-product H_2_O_2_ to prevent potential toxicity.

### Construction of recombinant *Bacillus subtilis* spores for intestinal nicotine degradation

To develop a catalytic machinery for intestinal application, we selected *Bacillus subtilis* spores to display the enzyme. Given that *B. subtilis* spores are inherently resistant to heat, acid, and proteolysis, oral administration of *B. subtilis* spores can pass through the gastrointestinal tract without persistent colonization (15), which is particularly important for fast onset and subsequent clearance. Therefore, we fused ssNox to eight spore coat proteins (CotB, CotC, CotE, CotG, CotX, CotY, CotZ, and OxdD) through genomic integration for the comparison as an anchor protein to facilitate surface display (Fig. 2A). Among the resulting recombinant strains, CotE-anchored construct, designated EN, exhibited the highest nicotine-degrading activity, and CotE was then chosen as the anchor protein (Fig. 2B, 2C; Fig. S5). To minimize spore germination in the intestine, which is potentially a factor to compromise catalytic performance, we deleted the germination-associated *gerKABC* operon in the genome. This modification markedly reduced spore germination while preserve dominant spore morphology after 24 h of incubation in simulated intestinal fluid (Fig. 2D).

**Figure 2.**
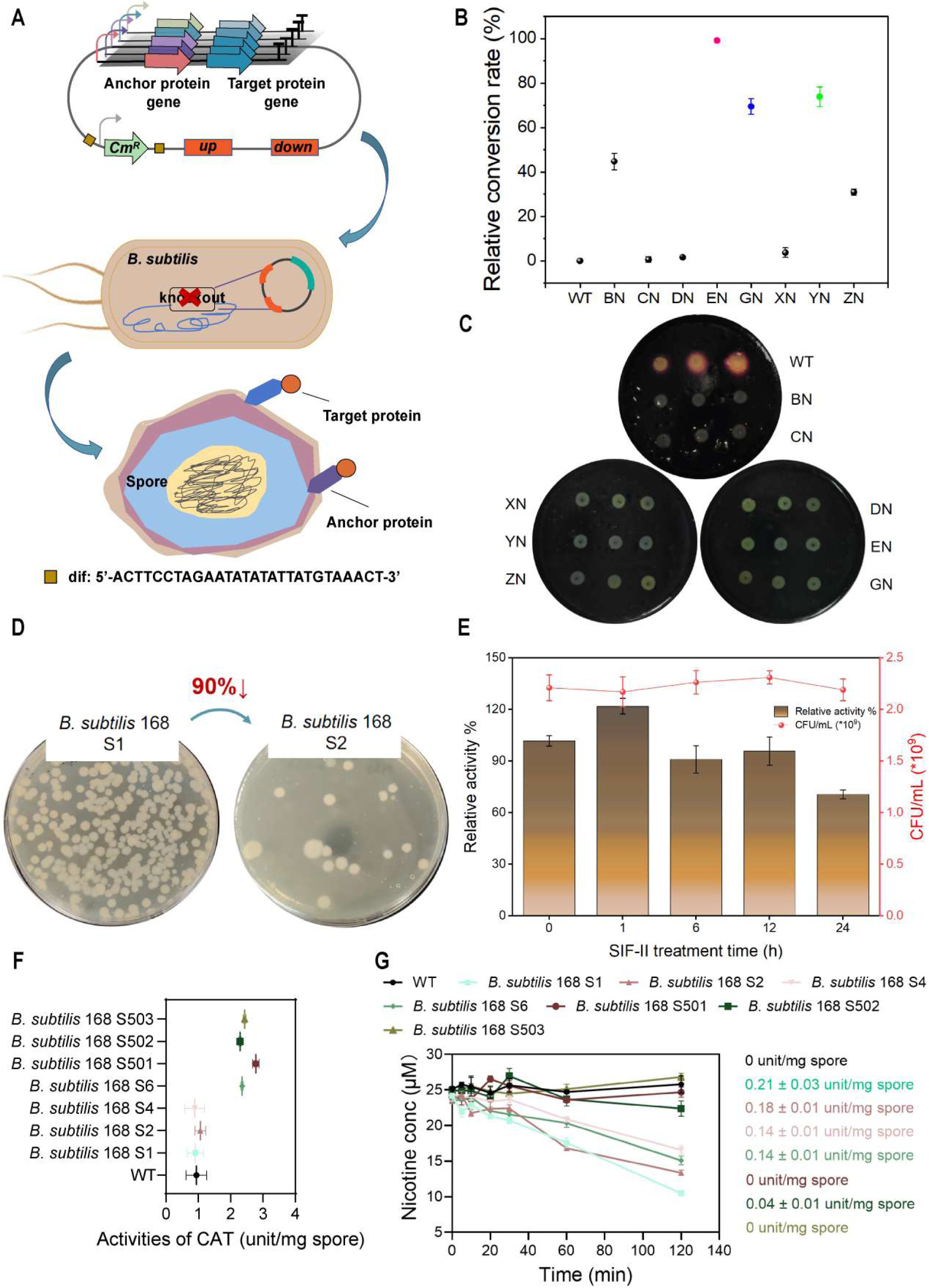
Surface display on *B. subtilis* spores for non-colonizing enzymatic degradation. (A) Illustration of genomic integration for displaying ssNox on spore surface. (B) Screening of spore coat anchor proteins based on relative nicotine-degrading activity. (C) Confirmation of genomic integration by insertional inactivation of amylase activity. The recombinant strains of BN, CN, DN, EN, GN, XN, YN, and ZN represent the fusions of ssNox to eight spore coat proteins of CotB, CotC, OxdD, CotE, CotG, CotX, CotY, and CotZ, respectively. (D) Verification of the germination-deficient phenotype. (E) Stability of recombinant spores in simulated intestinal fluid containing protease and bile salts (SIF-II). (F) Catalase activity of strains co-display ssNox and MnCAT. (G) ssNox activity of the recombinant strains, showing restoration of catalytic activity by dual-anchor approach. Data are presented as the mean ± SD (n = 3).

To decompose H_2_O_2_ generated during ssNox-catalyzed nicotine oxidation, we next incorporated a manganese catalase (MnCAT) derived from a thermophilic *Anoxybacillaceae* to prevent the accumulation of H_2_O_2_. Initial attempt by co-display of ssNox and MnCAT using the same CotE anchor protein resulted in the decrease of ssNox activity, possibly caused by steric interference between the displayed enzymes (Fig. 2G). We therefore adopted a dual-anchor approach, displaying MnCAT with CotE and ssNox with CotY, which restored the activities of both enzymes (Fig. 2F and G). To further improve the biosafety of the engineered strain, we precisely deleted the chloramphenicol-resistance marker using endogenous *Xer/dif* site-specific recombination system (16) (Fig. S7). The resulting strain, *B. subtilis* 168 S6, possessed both nicotine-oxidizing and H_2_O_2_-degrading activities without detectable effects on bacterial growth or sporulation under the tested conditions (Fig. S9). Through engineering and evaluation, a recombinant strain of *B. subtilis* 168 S6 was constructed, and its spores could be utilized to degrade nicotine and the by-product H_2_O_2_ in intestine.

### Encapsulation of enteric microspheres for intestinal release of ssNOX-displayed spores

Despite the trypsin resistance of engineered ssNox and the intrinsic stability of *B. subtilis* spores, oral administration requires additional protection from the damage by gastric acid and proteases. Oral delivery of macromolecules, particularly catalytically active enzymes, remains a challenging issue (17). To overcome this limitation, we fabricated pH-responsive enteric microspheres, designated rBS^EN^@SA&HA@CS MP, by combining ionic cross-linking with layer-by-layer assembly. The resulting core-shell structure comprised a calcium carbonate layer for acid neutralization, a hyaluronic acid-sodium alginate hydrogel network for structural support, and an outer chitosan layer for pH-responsive release. Changes in zeta potential after each coating step supported the success of sequential assembly for the preparation of the microspheres (Fig. 3B and Fig. S14).

**Figure 3.**
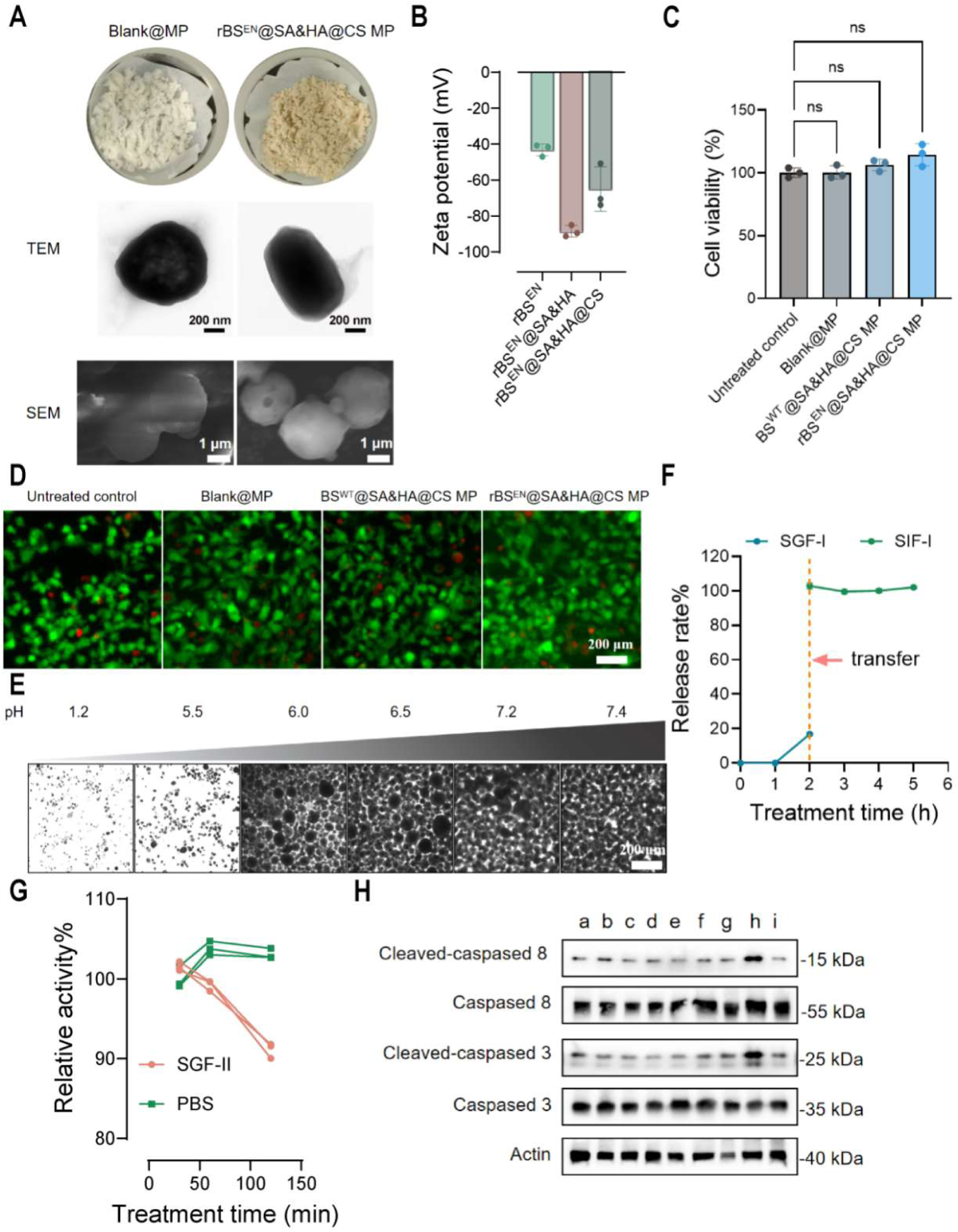
pH-responsive enteric microspheres for the protection of spore-displayed enzymes from gastric conditions and targeted intestinal release. (A) Morphological characterization of the microspheres by TEM and SEM. (B) Changes in zeta potential during sequential coating. (C) CCK-8 cell viability assay and (D) Calcein-AM/PI staining of IEC-6 cells exposed to the microspheres. (E) Swelling behaviors of the microspheres at different pH values. (F) Release of encapsulated spores during sequential incubation in simulated gastric fluid without pepsin (SGF-I) and simulated intestinal fluid without protease (SIF-I). (G) Residual nicotine-degrading activity after incubation in simulated gastric fluid containing pepsin (SGF-II). (H) Western blot analysis of apoptosis-associated proteins in IEC-6 cells, showing attenuation of H_2_O_2_-associated cytotoxicity by coupling ssNox–MnCAT. Lane a–i represent: PBS; Blank@MP; rBS^EN^@SA&HA@CS MP (*B. subtilis* S2); rBS^EN^@SA&HA@CS MP (*B. subtilis* S6); Blank@MP and 300 ng/mL nicotine; rBS^EN^@SA&HA@CS MP (*B. subtilis* S2) and 300 ng/mL nicotine; rBS^EN^@SA&HA@CS MP (*B. subtilis* S6) and 300 ng/mL nicotine; 300 ng/mL NIC and 0.1 mg/mL ssNox; rBS^EN^@SA&HA@CS MP (*B. subtilis* S6) and 300 ng/mL nicotine in the presence of 0.1 mg/mL ssNox.

The microspheres remained intact under acidic conditions at pH below 5.5, with a swelling ratio under 3, but rapidly swelled and disintegrated at pH greater than 6.0, consistent with the release profile in the intestinal environment (Fig. 3E; Fig. S15). In simulated gastrointestinal release assays, approximately 10% of the encapsulated spores were released during 2 h of incubation under gastric conditions, followed by sustained release after transferring to simulated intestinal fluid (Fig. 3F; Fig. S16). The intestinal-phase release profile appeared to be zero-order kinetics. After 2 h of incubation in simulated gastric fluid containing pepsin (SGF-II), encapsulated spores retained 91.1% of nicotine-degrading activity, whereas unencapsulated spores lost all activity within 30 min (Fig. 3G).

The cytocompatibility of the microspheres was evaluated in IEC-6 cells using CCK-8 and live/dead staining assays (Fig. 3C, 3D). Western blotting and flow cytometry further revealed that nicotine degradation by the microsphere-encapsulated system reduced H_2_O_2_-associated apoptotic signaling, represented by cleaved caspase-3/8 (Fig. 3H; Fig. S17). The proportion of apoptotic cells decreased from 33.8% to 8.1%, close to the blank. Taken together, the recombinant spores were encapsulated by multilayer sequential assembly with a satisfactory performance on the protection from gastric damage and targeted intestinal release. The coupling of ssNox and MnCAT on the recombinant spores also eliminated the toxic effects from H_2_O_2_ during nicotine degradation.

### Evaluation of oral administration of the microspheres on intestinal homeostasis

To ensure the full function of molecular siphoning by the microspheres, the duration time in the gastrointestinal tract could affect overall outcomes after oral administration, which may also interfere intestinal homeostasis to pose a safety concern. To address this, we therefore conducted a toxicity study with 14-day repeated-dosing by orally administering 15 mg of microspheres daily in mice (Fig. 4A). Mice in the microsphere-treated group showed a transient fluctuation in body weight during the dosing period (Fig. 4B), but no obvious abnormality was observed throughout the study. Serum analyses indicated no significant changes in either inflammatory markers, including IL-6 and TGF-β1, or the indices of hepatic and renal functions, including AST, ALT, TBIL, BUN and creatinine (Fig. 4C). Histological examination by H&E staining did not find detectable lesions, inflammatory cell infiltration, or structural abnormalities in the heart, liver, spleen, lungs, kidneys, brain, or intestine (Fig. 4D).

**Figure 4.**
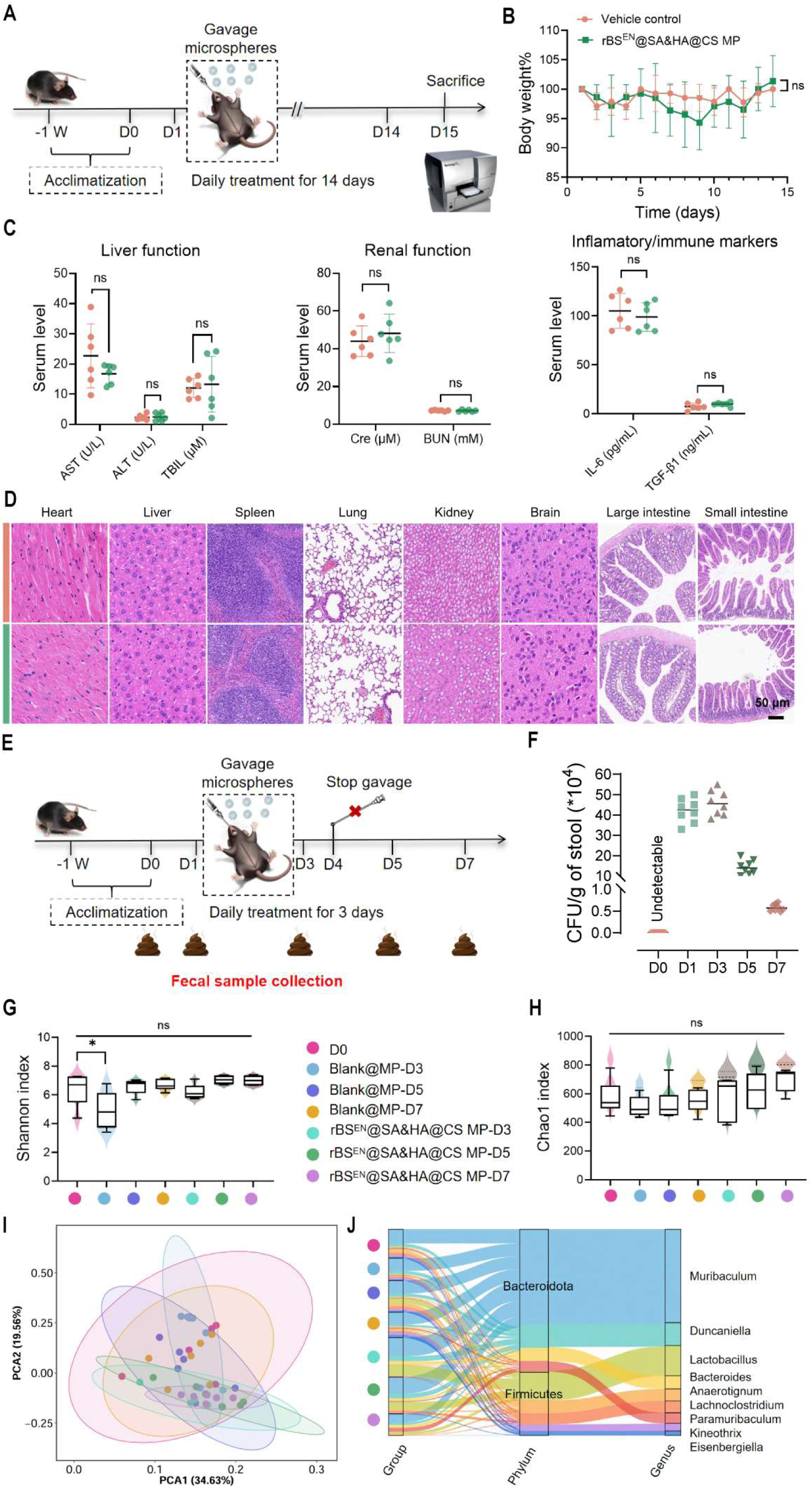
The effects of oral microspheres on gastrointestinal tract and gut homeostasis. (A) Experimental timeline for 14-d repeated-dose oral toxicity study. (B) Changes in body weight during microsphere administration. (C) Serum concentrations of IL-6 and TGF-β1 and biochemical indices of hepatic and renal functions. (D) Representative H&E stained sections of major organs. (E) Fecal sampling schedule for microbiome analysis. D0 indicates baseline samples collected immediately before the first oral gavage. (F) Viable spore counts in fecal samples during the treatment and washout periods (n = 8). (G) Chao1 richness and (H) Shannon diversity indices of the gut microbiota (n = 6). (I) PCA of gut microbial communities. (J) Sankey plot showing taxonomic composition across treatment groups. Data in (B) were analyzed by two-way ANOVA. Data in (G) and (H) were analyzed by ordinary one-way ANOVA. *P < 0.05; ns, not significant.

By viable counting the fecal samples, the spores administered in mice were progressively cleared in the feces, decreasing from 4.59 × 10^5^ CFU/g on day 3 to 0.58 × 10^5^ CFU/g on day 7 (Fig. 4F, S19A). Further analysis of the gut microbiota by 16S rRNA sequencing did not show significant differences in alpha or beta diversity among treatment groups (Fig. 4G–4I, S20–S21A). Although the relative abundance of the genus of *Lactobacillus* showed a trend of increase after the treatments (Fig. 4J), the probiotic nature of this genus was unlikely to relate pathological changes. Meanwhile, *B. subtilis* did not affect the relative abundances of 200 most abundant genera (Fig. S21B). Together, these data indicated that the orally administered system was largely cleared from the gastrointestinal tract after a week. The oral administration of the encapsulated microspheres did not produce either detectable systemic toxicity, or major changes of gut microbial community under the tested conditions.

### Evaluation of oral administration of the microspheres on central reward signaling

Next, we sought to determine whether a gut-localized enzymatic machinery could intercept nicotine in the gastrointestinal tract and thereby reduce its concentrations in the blood and brain following an acute nicotine challenge. Thus, we first evaluated the gastrointestinal transit of DiR-labeled microspheres in mice (Fig. S23A) and characterized the influence of nicotine dose on behaviors, aiming to establish an appropriate dosing interval and the dose for nicotine challenge. In vivo and ex vivo imaging showed that, after oral gavage for 1 h, the fluorescence signal was primarily concentrated in the stomach and duodenum; at 3 h, spreading throughout the small intestine and the beginning of accumulation in the cecum were observed (Fig. 5A, 5B). During a 24-h period, fluorescence signal was not able to detect in other tissues, including heart, liver, spleen, lung, kidney, or brain. In the open-field test, 0.5 mg/kg nicotine could induce noticeable locomotor activity, and 1.5 mg/kg produced marked hypolocomotion (Fig. S23C, S23D). As a result, we selected 0.5 mg/kg for subsequent evaluations. To examine the therapeutic efficacy, mice were orally treated with the microspheres 1 h before intravenous administration of nicotine to allow the microspheres locating in the small intestine. In the mouse model, 15 mg of rBS^EN^@SA&HA@CS MP resulted in a complete nicotine degradation within 1 h across the tested concentration range.

**Figure 5.**
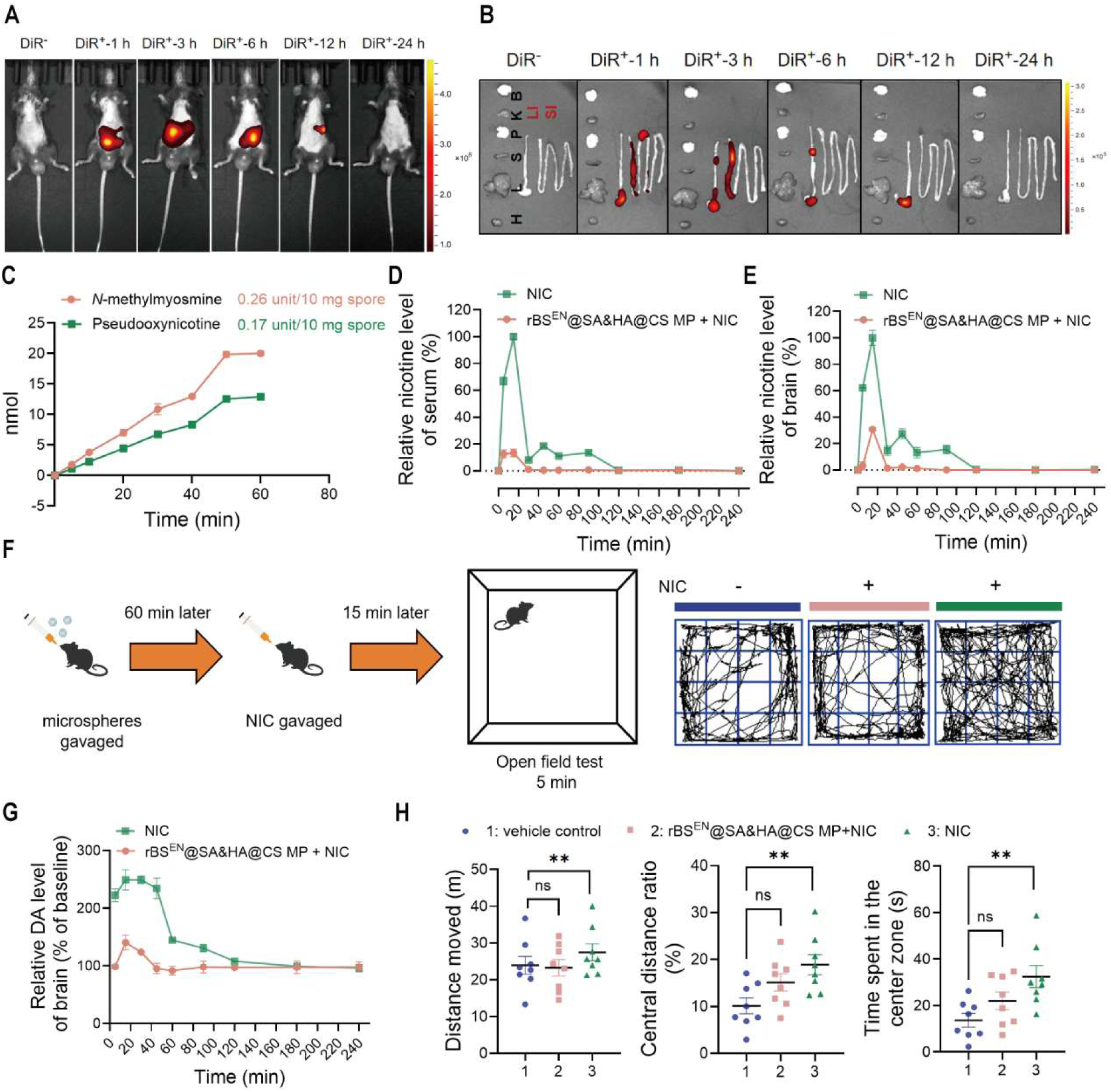
The effects of intestinal interception and in situ degradation of nicotine by the microspheres on central reward signaling. (A) In vivo and (B) ex vivo fluorescence imaging of DiR-labeled microspheres during gastrointestinal transit. (C) In vitro nicotine-degrading activity of the microspheres. (D) Blood and (E) brain nicotine profiles after nicotine administration with or without microsphere pretreatment (n = 3). (F) Experimental design and representative trajectories for the open-field test (OFT). (G) Brain dopamine concentrations after nicotine administration (n = 3). (H) Quantitative analysis of open-field behavior (n = 8). Data are presented as the mean ± SEM. Statistical comparisons were performed by one-way ANOVA followed by indicated multiple-comparisons test. *P < 0.05, **P < 0.01.

To assess the efficacy of the microspheres, further analyses were performed. In addition to the effective degradation of nicotine by the microspheres *in vitro* (Fig. 5C), microsphere pretreatment reduced the maximum nicotine concentrations in the brain and blood to 30.68 ± 1.20% and 13.35 ± 2.92% compared to those in the nicotine-only group (Fig. 5D, 5E). Consistently, microsphere pretreatment reduced nicotine-induced brain dopamine release from 249.35 ± 17.55% to 140.73 ± 12.57% judged from the baseline (Fig. 5G). Nicotine-induced hyperlocomotion was also attenuated from the pretreatment of the microspheres, as examined by open-field testing, which showed that the nicotine-induced increases in center-zone activity and total locomotor measures were both significantly reversed (Fig. 5F, 5H). Together, these findings confirmed that gut-localized enzymatic degradation could reduce nicotine levels in the blood and brain and attenuate nicotine-associated dopaminergic and behavioral responses.

### Examination of oral administration of the microspheres on addiction-related neuroadaptation and systemic toxicity

To evaluate whether intestinal nicotine degradation driven by molecular siphoning could modulate addiction-related phenotypes during chronic nicotine exposure, a spontaneous withdrawal model was employed (Fig. 6A) to assess withdrawal-associated hyperalgesia by measuring mechanical paw withdrawal threshold (PWT). During the period of nicotine exposure, mice in nicotine-only group at Day 10 exhibited a reduction of PWT from 1.0 g at the baseline to 0.6 g, whereas mice treated with the microspheres maintained PWT values close to the baseline (Fig. 6B). Following dose escalation (Day 14), PWT remained higher in the treatment group than that in nicotine-only group (0.6 vs. 0.4 g). After spontaneous withdrawal, mice in nicotine-only group showed lower locomotor activity, with a decrease of average speed by 44.2% and a decline of total travel distance from 24.89 ± 4.76 m at the beginning to 13.73 ± 3.23 m, together with anxiety-associated behaviors, such as whisker loss (Fig. 6C, S25). Both preventive and therapeutic administration of the microspheres restored locomotor performance to the levels comparable to the baseline.

**Figure 6.**
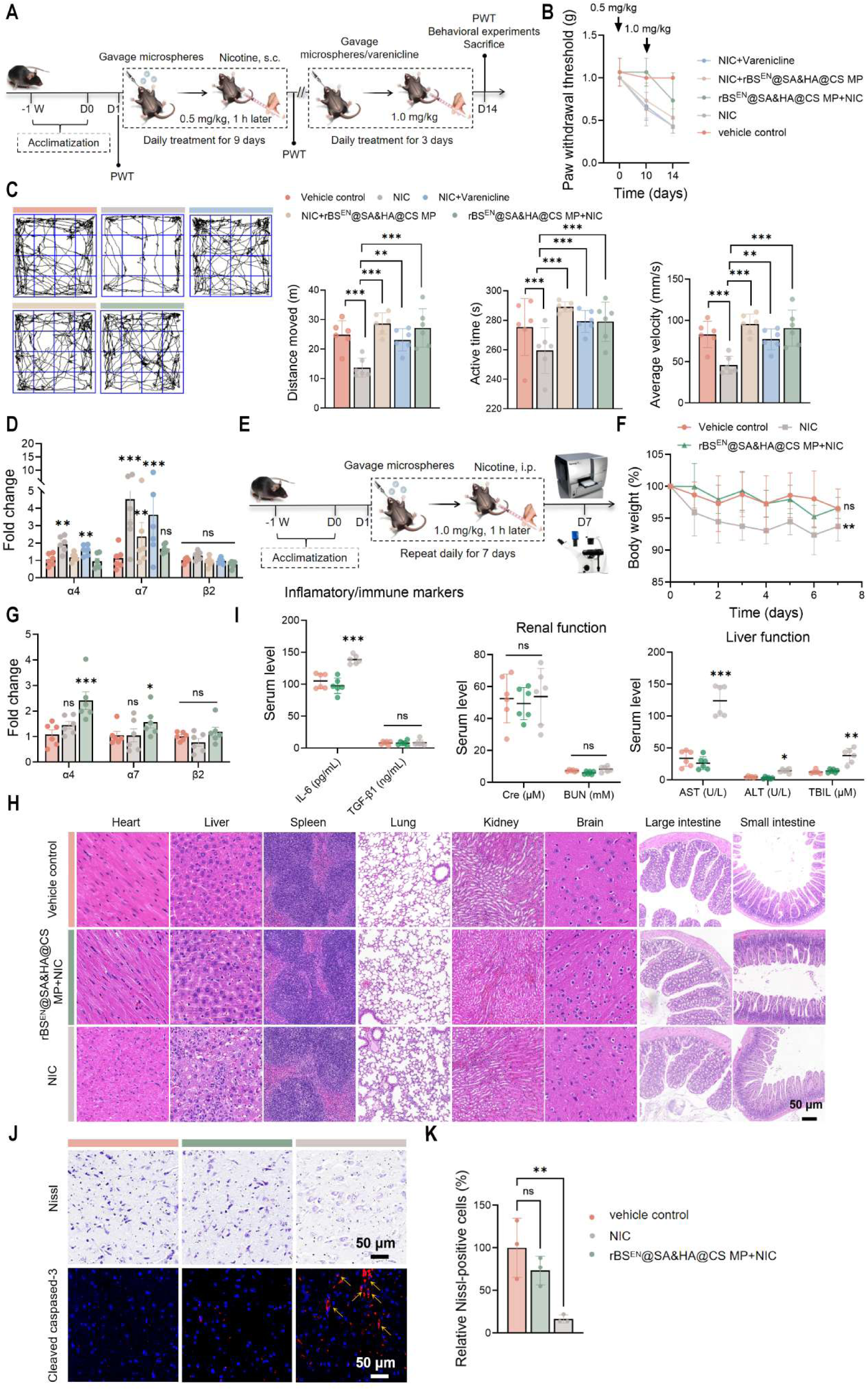
Addiction-related neuroadaptation and systemic toxicity by the treatments. (A) Experimental design for the chronic nicotine exposure and spontaneous withdrawal model. (B) Mechanical paw withdrawal thresholds during nicotine exposure and withdrawal. (C) OFT behaviors following spontaneous withdrawal. (D) Relative expression of α4, α7, and β2 nAChR subunits in whole-brain samples from chronic nicotine exposure model. (E) Experimental timeline for the high-dose nicotine toxic model. (F) Body weight trajectories during nicotine administration. (G) Relative expression of nicotinic acetylcholine receptor subunits in whole-brain samples from the toxic model. (H) Representative H&E-stained sections of major organs. (I) Serum biochemical indices of hepatic and renal functions. (J) Representative Nissl staining and cleaved caspase-3 immunofluorescence of brain sections. (K) Quantification of the relative content of Nissl-positive cells. *P < 0.05, **P < 0.01, ***P < 0.001 compared with vehicle control or NIC group.

Because chronic nicotine exposure has been associated with increased expression of nicotinic acetylcholine receptors (nAChRs) (18), we next quantified the levels of receptor expression by qPCR. The expressions of α4 and α7 nAChR subunits were significantly increased in nicotine-only group (Fig. 6D). Preventive microsphere treatments prevented the increase, maintaining receptor expression unchanged. In contrast, varenicline, the partial agonist of nicotine receptor (19), could alleviate withdrawal-associated behavioral changes, but did not prevent the elevation of nAChR expression (Fig. 6D).

Next, to evaluate the protective effects of intestinal nicotine degradation at high dose, a toxic nicotine model (1.0 mg/kg, intraperitoneally, once daily for 7 days), corresponding to nicotine exposure previously estimated as the amount in heavy smokers (20), was used for the examination. When the mice were pretreated with the microspheres, nicotine-associated body weight loss was not observed (Fig. 6F). In addition, serum markers of liver injury, including AST, ALT and TBIL, which were approximately 3.0- to 3.6-fold higher in the nicotine group, were in the normal range (Fig. 6I). Histological examination further revealed the reduction of hepatic steatosis, inflammatory infiltration and pulmonary interstitial injury by microsphere treatment (Fig. 6H). Nissl staining and immunofluorescence analysis of cleaved caspase-3 additionally indicated preservation of neuronal morphology and reduced apoptosis in the brain (Fig. 6J, 6K). Together, these findings indicated that molecular siphoning-driven intestinal nicotine degradation attenuated addiction-related behavioral and molecular adaptations, and significantly reduced the systemic toxicity associated with chronic nicotine exposure.

## Discussion

In this study, we developed an orally administered, gut-restricted enzymatic platform to reduce systemic nicotine exposure through intestinal molecular siphoning. By combining an engineered nicotine-degrading enzyme, surface display on *B. subtilis* spores and multilayer encapsulation, the resulting microspheres retained catalytic activity under gastrointestinal conditions and reduced nicotine concentrations in both blood and brain. Oral administration also attenuated nicotine-induced dopamine release, normalized the expression of α4 and α7 nicotinic acetylcholine receptors, alleviated withdrawal-associated hyperalgesia, locomotor impairment and anxiety-related behaviors, and reduced nicotine-associated injury in liver, lung, and brain. Together, these findings demonstrate that local degradation of a circulating small molecule in the intestinal lumen can alter its systemic and neurobiological effects without requiring systemic exposure of the therapeutic agent. Meanwhile, the present results support the dosing rationale for the enzymatic degradation at nicotine concentrations relevant to human exposure (3), despite that intestinal pharmacokinetics and dosing escalations would be required in humans.

The effects of intestinal molecular siphoning predominately depend on the efficiency of sustained degradation of nicotine in the intestinal lumen. Therefore, systemic clearance of nicotine not only correlates to intestinal permeability, but also the capacity and persistence of the luminal catalytic machinery, in which the real-time kinetic analysis is valuable. However, owing to the complexity of intestinal contents to interfere with quantitative detections, transintestinal nicotine flux was not able to be directly measured in this study, requiring further exploration. In principle, intestinal molecular siphoning differs from any existing pharmacological and enzymatic approaches to treat nicotine dependence. Currently, nicotine replacement therapy reduces withdrawal symptoms by maintaining lower but persistent systemic nicotine exposure and therefore continues to position the neuronal system to the addictive molecule (21). As a partial agonist for nAChR, Varenicline can reduce craving and withdrawal, but it is associated with various psychiatric adverse events (6). Systemically admonitions of nicotine-degrading enzymes can lower blood nicotine concentrations, but repeated intravenous administration and strong immunogenicity prevent them from clinically useful therapeutics (7, 9). Differently, the present system localizes the catalytic machinery in the intestinal lumen, thereby avoiding potential adverse events from systemic exposures. Meanwhile, the spore-display system also provides a rapid onset without the need of stable intestinal colonization, distinguishing it from other live biotherapeutic strategies that depend upon bacterial engraftment or sustained metabolic adaptation. Given that high-dose nicotine administration also increases oxidative stress, as indicated by elevated reactive oxygen species and lipid peroxidation in brain (22, 23), the present microsphere treatments mitigate these effects and reduce histological injury in multiple organs. Moreover, nicotine toxicity is reflected by the combination of concentration, exposure duration, and cell type (24–26), and chronic exposure to nicotine-containing products at low nicotine concentration can also cause persistent injury to the cardiovascular endothelium (27). Therefore, the present therapeutic rationale could extend the context beyond preventing acute cellular toxicity to the reduction of cumulative systemic consequences of nicotine exposure.

Safety is a pivotal factor for any therapeutic agents, which usually is a major hurdle in drug discovery and development. In our case, the targeted release for localized function and the independence of specific targets can theoretically avoid systemic exposure, broad tissue distribution and long residence time, exhibiting obvious advantages on the safety to a body. In this study, no microsphere-associated fluorescence was detected during 24-h period after oral administration of DiR-labeled microspheres, indicating an intestine-restricted distribution. Also, fecal viable counts showed a progressive elimination of the engineered spores, suggesting a transient gastrointestinal passage rather than persistent colonization, which agrees with a previous report (15). Furthermore, repeated oral dosing for 14 d did not produced any abnormality in serum biochemical indices or major-organ histology. Regarding the impact on gut microbiota by the microspheres, the relative abundance of *Lactobacillus* was marginally increased after the treatments. Given the probiotic nature of *Lactobacillus* genus, the biological significance of this change might be beneficial to the body. Overall, although long-term safety studies are required before clinical translation, the present data in animal models have strongly supported the safety of this therapeutic approach.

As a major implication of this work, intestinal molecular siphoning could be adaptable to other molecules beyond nicotine. Principally, this strategy could be applied to circulating small molecules that can cross the intestinal barrier and for which an efficient degrading enzyme is available or can be engineered. Thus, potential applications may include exogenous toxins, endogenous metabolites, and addictive substances. However, the feasibility of each target molecule would depend on several factors, including its physicochemical properties, bidirectional intestinal permeability, systemic production rate, concentration range, and the catalytic efficiency and the specificity of corresponding enzyme (s). In this aspect, the modularity of the spore-display platform may facilitate substitution of the catalytic component while retain the same oral delivery architecture. Nevertheless, several limitations of this study should be considered. First, the neurochemical analyses were performed using whole-brain samples and therefore cannot precisely represent region-specific effects in areas involved in reinforcement and withdrawal. Future studies should incorporate region-specific neurochemical measurements, spatial transcriptomic analyses, or real-time monitoring of nicotine and dopamine dynamics. Second, although the spontaneous withdrawal model captured somatic and behavioral features of dependence, it did not directly assess voluntary nicotine intake, craving or relapse. Therefore, self-administration, reinstatement, and long-term abstinence models are more favorable on evaluating whether intestinal nicotine degradation reduces motivational aspects of addiction. Third, the present therapeutic agent relies on enteric microspheres for gastric protection and intestinal release. Further enzyme engineering, such as introducing acid-resistant mutations through molecular design, may simplify the form and process while preserve catalytic activity. In addition, given the differences in nicotine sensitivity and metabolism across animal species and humans (28, 29), systemic nicotine exposure and the magnitude of benefits achieved through intestinal degradation are difficult to predict. In humans, variability in nicotine-metabolizing enzymes, such as CYP2A6, may similarly affect treatment responses and dosing requirements. Moreover, nicotine dependence is influenced by social context, stress, and learned behaviors (30, 31), which is unlikely to be addressed by metabolic intervention alone. The strategy of intestinal molecular siphoning would therefore be expected to complement, rather than replace, behavioral counseling and psychosocial support. In conclusion, this study demonstrates that gut-confined enzymatic degradation can substantially reduce systemic and cerebral nicotine exposure and attenuate associated neurochemical, behavioral, and toxicological effects in mice. These findings establish a proof of concept of intestinal molecular siphoning as a new avenue of therapeutic applications and provide a foundation for developing orally administered degradation therapies that remove circulating small molecules without requiring systemic delivery of the therapeutic agents.

## Materials and Methods

### Strains, plasmids, and culture conditions

The strains and plasmids used in this study are listed in Table S2. *E. coli* DH5α was used for plasmid construction, and *E. coli* BL21(DE3) was used for recombinant protein expression. *B. subtilis* WB800N and *B. subtilis* 168 were used as hosts for construction of the engineered strains. Detailed procedures for strain cultivation, plasmid construction, and genetic manipulation are provided in *SI Appendix* or was described previously (32).

### Animal studies

Male C57BL/6 mice (6–8 weeks, 22–26 g) were obtained from Hangzhou Ziyuan Laboratory Animal Technology Co., Ltd. Mice were housed under SPF conditions at 20–22 ℃ and 55 ± 5% relative humidity on a 12-h light/dark cycle, with ad libitum access to food and water. All animal care and procedures were approved by the Animal Ethics Committee of China Pharmaceutical University (approval no. YSL-202505037). For the repeated-dose oral toxicity study, mice received 15 mg of microspheres once daily by oral gavage for 14 d. The gastrointestinal distribution of DiR-labeled microspheres was monitored using an IVIS Lumina III in vivo imaging system. For the chronic nicotine exposure model, mice received repeated subcutaneous injections of nicotine for 9 d at 0.5 mg/kg, followed by dose escalation to 1.0 mg/kg for 3 d. Spontaneous withdrawal-associated behaviors were assessed after nicotine administration was discontinued. For the high-dose nicotine toxic model, mice received nicotine intraperitoneally at 1.0 mg kg once daily for 7 d. Microspheres were suspended in 0.5% (w/v) sodium carboxymethyl cellulose before oral administration. Nicotine solutions for oral gavage were prepared in 0.1% (w/v) sodium saccharin, while those for injection were prepared in saline, and both were freshly made. Sodium saccharin at this concentration was previously reported not to confound the behavioral outcomes evaluated in this study (33, 34).

### Statistical analysis

Data are presented as mean ± SD or mean ± SEM, as specified in the figure legends. Comparisons between two groups were performed using unpaired two-tailed Student’s t-test. Comparisons among three or more groups were performed using ordinary one-way ANOVA followed by Tukey’s or Dunnett’s multiple-comparisons test, as appropriate. Two-way ANOVA was used for experiments involving two independent factors. A value of P < 0.05 was considered statistically significant. ns denotes no significant difference. Statistical analyses were performed using GraphPad Prism 10 and Origin 2025b. Detail Materials and Methods are provided in *SI Appendix*.

## Acknowledgments

This work was supported by the grants from the National Key Research and Development Program of China (2018YFA0902000 to Y.C), the Key Research and Development Project of Guangdong Province (2022B1111070004 to Y.C.), and the Project Program of the State Key Laboratory of Natural Medicines, China Pharmaceutical University (SKLNMZZ202606 to Y.C).

## Data Availability Statement

The original 16S rRNA sequencing data are available from the Genome Sequence Archive (GSA) at https://ngdc.cncb.ac.cn/ under accession number PRJCA062447. Other data are available from the corresponding author upon reasonable request.

## Author Contributions

Y. C. conceived, designed and supervised the research, and acquired funding; X. X., R. Z., and Y. H. designed and performed research, analyzed data, and interpreted results; X. X. and Y. C. wrote the paper; M. Y., X. X., and Y. S. performed research. All authors approved the final version of the manuscript. Competing Interest Statement: Two Chinese patent applications (CN202511773173.1, CN2025114332212.2) were filed by China Pharmaceutical University, and Y. C., X. X., R. Z. and Y. H. were listed as inventors of the patents. Other authors declare no conflicts of interest.

## References

1. Le FB et al. Tobacco and nicotine use. Nat Rev Dis Primers 8, 19 (2022).

2. US Department of Health and Human Services. The Health Consequences of Smoking—50 Years of Progress: A Report of the Surgeon General (US Department of Health and Human Services, CDC, 2014). Available at: https://www.ncbi.nlm.nih.gov/books/NBK179276

3. Chen B et al. Gut bacteria alleviate smoking-related NASH by degrading gut nicotine. Nature 610, 562–568 (2022).

4. IHME. Global Burden of Disease (2025): Burden by Risk, 1990–2023 (Institute for Health Metrics and Evaluation, 2025). Available at: https://ourworldindata.org/grapher/number-of-deaths-by-risk-factor

5. Benowitz NL. Nicotine addiction. N Engl J Med 362, 2295–2303 (2010).

6. Bagdas D et al. New insights on the effects of varenicline on nicotine reward, withdrawal and hyperalgesia in mice. Neuropharmacology 138, 72–79 (2018).

7. Xue S, Schlosburg JE, Janda KD. A new strategy for smoking cessation: characterization of a bacterial enzyme for the degradation of nicotine. J Am Chem Soc 137, 10136–10139 (2015).

8. Pentel PR et al. The nicotine-degrading enzyme NicA2 reduces nicotine levels in blood, nicotine distribution to brain, and nicotine discrimination and reinforcement in rats. BMC Biotechnol 18, 46 (2018).

9. Xue S et al. An enzymatic advance in nicotine cessation therapy. Chem Commun (Camb) 54, 1686– 1689 (2018).

10. Kallupi M, Xue S, Zhou B, Janda KD, George O. An enzymatic approach reverses nicotine dependence, decreases compulsive-like intake, and prevents relapse. Sci Adv 4, eaat4751 (2018).

11. Thisted T et al. Optimization of a nicotine degrading enzyme for potential use in treatment of nicotine addiction. BMC Biotechnol 19, 56 (2019).

12. Abramson J et al. Accurate structure prediction of biomolecular interactions with AlphaFold 3. Nature 630, 493–500 (2024).

13. Siepen JA, Keevil EJ, Knight D, Hubbard SJ. Prediction of missed cleavage sites in tryptic peptides aids protein identification in proteomics. J Proteome Res 6, 399–408 (2007).

14. Gershon PD. Cleaved and missed sites for trypsin, lys-C, and lys-N can be predicted with high confidence on the basis of sequence context. J Proteome Res 13, 702–709 (2014).

15. Yang M et al. Engineered *Bacillus subtilis* as oral probiotics to enhance clearance of blood lactate. ACS Synth Biol 14, 101–112 (2025).

16. Bloor AE, Cranenburgh RM. An efficient method of selectable marker gene excision by Xer recombination for gene replacement in bacterial chromosomes. Appl Environ Microbiol 72, 2520– 2525 (2006).

17. Drucker DJ. Advances in oral peptide therapeutics. Nat Rev Drug Discov 19, 277–289 (2020).

18. Han P et al. Pharmacokinetic differences in nicotine and nicotine salts mediate reinforcement-related behavior: an animal model study. Front Neurosci 17, 1288102 (2023).

19. Marks MJ et al. Chronic treatment with varenicline changes expression of four nAChR binding sites in mice. Neuropharmacology 99, 142–155 (2015).

20. Kershbaum A, Bellet S, Khorsandian R. Elevation of serum cholesterol after administration of nicotine. Am Heart J 69, 206–210 (1965).

21. J Hartmann-Boyce, SC Chepkin, W Ye, C Bullen, T Lancaster, Nicotine replacement therapy versus control for smoking cessation. Cochrane Database Syst Rev 5, CD000146 (2018).

22. Newman MB et al. Nicotine’s oxidative and antioxidant properties in CNS. Life Sci 71, 2807–2820 (2002).

23. Khallaf MF, El-Bakary NM, El-Sayed IH, El-Sayed IH. Dose-related effects of nicotine on oxidative injury in young, adult and old rats. J Environ Biol 33, 233–238 (2012).

24. Li J et al. Cell death induced by nicotine in human neuroblastoma SH-SY5Y cells is mainly attributed to cytoplasmic vacuolation originating from the trans-Golgi network. Food Chem Toxicol 185, 114431 (2024).

25. Kim CS et al. Nicotine-induced apoptosis in human renal proximal tubular epithelial cells. PLoS One 11, e0152591 (2016).

26. Chioran D et al. Nicotine exerts cytotoxic effects in a panel of healthy cell lines and strong irritating potential on blood vessels. Int J Environ Res Public Health 19, 8881 (2022).

27. Münzel T, Crea F, Rajagopalan S, Lüscher TF. Nicotine and the cardiovascular system: Unmasking a global public health threat. Eur Heart J 47, 1764–1781 (2025).

28. Akinola LS et al. C57BL/6 substrain differences in pharmacological effects after acute and repeated nicotine administration. Brain Sci 9, 244 (2019).

29. Seemiller LR et al. Genetic differences in nicotine sensitivity and metabolism in C57BL/6J and NOD/ShiLtJ mouse strains. Neuropharmacology 221, 109279 (2022).

30. Christakis NA, Fowler JH. The collective dynamics of smoking in a large social network. N Engl J Med 358, 2249–2258 (2008).

31. Kassel JD, Stroud LR, Paronis CA. Smoking, stress, and negative affect: correlation, causation, and contextual mechanisms. Psychol Bull 129, 270–304 (2003).

32. Xiong X et al. Engineering *Bacillus* spores to display nicotine oxidase: *in situ* specific and sensitive nicotine detection. ACS Sens 10, 3589–3599 (2025).

33. Thompson Z, Kolb EM, Garland T Jr. High-runner mice have reduced incentive salience for a sweet-taste reward when housed with wheel access. Behav Processes 146, 46–53 (2018).

34. Russo M, Funk D, Loughlin A, Coen K, Lê AD. Effects of alcohol dependence on discrete choice between alcohol and saccharin. Neuropsychopharmacology 43, 1859–1866 (2018).

