## Supplementary Tables and Figures for "Intestinal molecular siphoning by engineered bacterial spores for non-systemic treatment of nicotine addiction"

**Yijun Chen*

*.*


**This file includes:**

Supporting Materials and Methods

Figures S1 to S27

Tables S1 to S3

Supporting Sequences

SI References

#### Supporting Materials and Methods

##### 1. Mining and engineering of nicotine-degrading enzyme ssNox

*1.1. Homology search and phylogenetic analysis*

NicA2 (WP_013973880.1) from *Pseudomonas putida* S16 was used as the query sequence for BLASTP searches against the NCBI non-redundant protein database. The 500 bacterial protein sequences with the highest similarity to NicA2 were selected for phylogenetic analysis. A phylogenetic tree was constructed using TBtools-II (1) and visualized via the iTOL. Proteins located within the same phylogenetic tree were performed using MAFFT. The three candidate proteins with the highest sequence similarity to NicA2 were aligned again, and the alignment was visualized using ESPript 3.0.

*1.2. Structural and functional prediction and screening*

Potential protease cleavage sites in the candidate proteins were predicted using PeptideCutter and further evaluated according to the Keil rules (Table S1) (2, 3). Surface solvent accessibility was analyzed using I-TASSER and NetSurfP-3.0. These predictions were interpreted together with three-dimensional structural models generated using AlphaFold 3 (4). A previously uncharacterized NicA2 homolog containing the fewest predicted trypsin cleavage sites was selected for subsequent engineering.

*1.3. Semi-rational design and mutant library construction*

Previous studies have shown that reducing surface hydrophobicity or increasing positive surface charge can improve protein thermal stability and resistance to pepsin and trypsin (5). On the basis of the sequence and structural analyses described above, residues at the most probable trypsin cleavage sites were selected for targeted mutagenesis. Because improved thermal stability is often associated with increased resistance to proteolysis, potential mutation sites for enhancing ssNox thermostability were also predicted using HotSpot Wizard. Mutations predicted to improve protease resistance or thermal stability were combined to construct the mutant library. Mutations were introduced by overlap-extension PCR. Protein expression, purification, and mutagenesis were performed as described previously (6).

*1.4. Protease tolerance assays*

The proteolytic tolerance of WT ssNox was first evaluated using pepsin and trypsin. Pepsin (≥ 2500 U/mg, Sangon Biotech, Shanghai, China) and trypsin (≥ 250 U/mg, Sangon Biotech) were prepared in 0.1 M Gly–HCl buffer, pH 2.0, and 0.1 M Tris–HCl buffers, pH 7.5, respectively. Protease solutions at 1 mg/mL were mixed with purified ssNox at a protease-to-protein mass ratio of 1:50. The mixtures were incubated at 37 °C with shaking at 300 rpm for 4 h. Residual nicotine-degrading activity was then measured at 37 °C and pH 7.5 for 30 min and quantified by LC-MS/MS. For the evaluation of trypsin tolerance in ssNox mutants, purified protein and trypsin were mixed at a trypsin-to-protein mass ratio of 200:1 and incubated at 37 °C for the indicated durations. Proteolysis was terminated by adding an equal volume of protease inhibitor solution. One aliquot was used to determine residual enzymatic activity, and the remaining sample was analyzed by SDS-PAGE.

#### 2. LC-MS/MS quantification

Analytes were quantified using a Shimadzu LCMS-8045 triple-quadrupole mass spectrometer operated in positive electrospray ionization mode with multiple-reaction monitoring. The monitored transitions were m/z 163.15>130.15 for nicotine (NIC), 161.10>119.10 for *N*-methylmyosmine (NMM), 179.01>106.15 for pseudooxynicotine (PON), and 166.01>130.15 for nicotine-*d3* (NIC-*d3*), which was used as the internal standard (Table S3).

Chromatographic analysis was performed using a C18 column with 95% methanol as the mobile phase at 30 °C. Steady-state kinetic parameters were determined by measuring initial reaction velocities over a nicotine concentration range of 1.0 to 800 μM. The resulting data were fitted to the Michaelis–Menten equation to obtain $K_m$ and $V_{max}$, and $k_{cat}$ was calculated from the enzyme concentration used in the assay. NIC was purchased from Chengdu Lemeitian Pharmaceutical Technology Co., Ltd. NIC-*d3* was obtained from Shanghai Zhenzhun Biological Technology Co., Ltd.

#### 3. Construction and characterization of recombinant spores

##### *3.1. Screening of spore coat anchor proteins*

*B. subtilis* WB800N was initially used to screen candidate spore coat anchor proteins. This strain lacks eight extracellular proteases encoded by *nprE*, *nprB*, *aprE*, *epr*, *mpr*, *bpr*, *vpr*, and *wprA* (7), thereby reducing extracellular proteolysis of heterologously expressed proteins. The coding sequences of the spore coat proteins CotB, CotC, OxdD, CotE, CotG, CotX, CotY, and CotZ, together with their native promoters, were amplified from the *B. subtilis* genome using the corresponding primer pairs. Each amplified fragment was assembled with the ssNox coding sequence and cloned into pDG364 by seamless cloning. Recombinant plasmids were verified by restriction-enzyme digestion and DNA sequencing performed by Sangon Biotech, Tsingke Biotechnology, or Gencefe Biotech. The verified plasmids were transformed into competent *B. subtilis* WB800N cells. Chromosomal integration at the *amyE* locus was confirmed by genomic PCR using primers amyE-F/R (5′-

CTTTTTTGTTTGGAAAGCGAGGGAAG-3′/5′-GGTTTAAATCCATCACATCGCCCATTG-3′) with an annealing temperature of 56.8 °C. Amylase activity assays, spore preparation, extraction of spore coat proteins, Western blot analysis, and nicotine-degradation assays were performed as described previously (6). The resulting recombinant strains displaying ssNox through CotB, CotC, OxdD, CotE, CotG, CotX, CotY, or CotZ were designated BN, CN, DN, EN, GN, XN, YN, and ZN, respectively.

##### *3.2. Construction of the B. subtilis 168-derived strains*

*B. subtilis* 168 was used as the chassis strain for subsequent strain construction. This strain is auxotrophic for tryptophan and does not contain additional antibiotic-resistance determinants. The overall construction strategy is shown in Fig. S6. The DNA fragment encoding the selected ssNox mutant was fused to CotE and cloned into pDG364 to generate pDG364-01. The resulting expression cassette was integrated into the chromosomal *amyE* locus of *B. subtilis* 168, yielding strain S1.

To reduce germination of the recombinant spores in the intestinal environment, the *gerKABC* operon was deleted. The upstream and downstream homology arms flanking *amyE* in pDG364-01 were replaced with sequences flanking the *gerKABC* operon, generating pDG364-02. Integration of this construct into the *B. subtilis* 168 chromosome produced the germination-deficient strain S2.

The chloramphenicol-resistance gene (*Cmr*) was used as the selection marker during strain construction. To remove this marker, the endogenous Xer/dif site-specific recombination system was used. Xer is a site-specific recombinase in *B. subtilis* that recognizes dif sites and mediates recombination between two identical sequences, thereby excising the intervening DNA fragment (9). A dif sequence (5′-ACTTCCTAGAATATATATTATGTAAACT-3′) was introduced on each side of the *Cmr* cassette in pDG364-02 by PCR, generating pDG364-03. Integration of this construct produced strain S3, from which the resistance cassette could be excised by Xer-mediated recombination. To increase expression of the surface-displayed enzyme, the native cotE promoter in pDG364-03 was replaced with the constitutive P43 promoter amplified from the *B. subtilis* 168 genome. The resulting plasmid, pDG364-04, was used to construct strain S4.

##### *3.3. Construction of ssNox and MnCAT co-display strains*

The MnCAT coding sequence, derived from a thermophilic member of the family *Anoxybacillaceae*, was synthesized by MiaoLing Bio Co., Ltd. MnCAT was fused to the *CotE-ssNox* display cassette through peptide linkers of different lengths. The corresponding constructs were introduced into *B. subtilis* 168 to generate strains S501, S502, and S503. To reduce interference between the co-displayed enzymes, strain S6 was constructed using pDG364-06, which contained separate CotY-ssNox and CotE-MnCAT expression cassettes. Chromosomal modification of the resulting strains was verified by genomic PCR using primers gerKABC-F/R(5′-TGCAGACAATATCAGCATCCGACCCATTGGTACAGGATGAGTCTATC-3′/5′-GGCGCACCCTATATCCCCCTTT-3′) with an annealing temperature of 57.5 °C. Genomic DNA was isolated using a bacterial genomic DNA extraction kit (Accurate Biology) according to the manufacturer's instructions.

#### 4. Evaluation of $H_2O_2$-associated cytotoxicity during ssNox-mediated nicotine degradation

IEC-6 cells were used for evaluate cytotoxicity by CCK-8 assay, Calcein-AM/PI live/dead staining, flow cytometry for apoptosis analysis, and Western blot analysis of cleaved caspase-3/8. Cytotoxicity associated with $H_2O_2$ generated during ssNox-mediated nicotine degradation was assessed by incubating IEC-6 cells with ssNox and nicotine in the presence or absence of catalase (CAT). The experimental groups were as follows: (a) negative control (PBS); (b–d) nicotine (NIC) at 3, 30, and 300 ng/mL; (e–g) NIC (3, 30, and 300 ng/mL, respectively) combined with 0.1 mg/mL ssNox; (h) 300 ng/mL NIC + 0.1 mg/mL ssNox + 5 μg/mL CAT; (i) 300 ng/mL NIC + 5 μg/mL CAT; (j) 0.1 mg/mL ssNox alone; (k) 5 μg/mL CAT alone; and (l) positive control (100 μM $H_2O_2$). After 24 h of incubation, cellular morphology was examined by microscopy. Cell viability was measured using the SuperKine™ Maximum Sensitivity Cell Counting Kit-8 (Abbkine Scientific Co., Ltd.) and assessed by Calcein-AM/PI Live/Dead Double Staining Kit (Yeasen Biotech). Apoptosis and necrosis were quantified by flow cytometry using a Cell Apoptosis and Necrosis Detection Kit (Beyotime Biotechnology). For Western blot analysis, total cellular protein was extracted after treatment and subjected to SDS-PAGE and membrane transfer. The membranes were probed with primary antibodies against cleaved caspase-3 and cleaved caspase-8 purchased from ImmunoWay Biotechnology. An antibody against β-actin purchased from ABclonal Biotechnology Co., Ltd. was used as the loading control. Protein signals were detected and quantified relative to β-actin. All commercial kits were used according to the manufacturers' instructions.

#### 5. Assessment of biological activities of recombinant spores

##### *5.1. Nicotine-degrading activity assay*

Recombinant spores were harvested from cultures grown in DSM for 96 h by centrifugation, washed with PBS, and adjusted to an $OD_{600}$ of 1.0. A 200-μL aliquot of the spore suspension was mixed with 2 μL of 5 mM nicotine stock solution and incubated at 37 °C. At 0, 5, 10, 20, 30, 60, and 120 min, an aliquot of the reaction mixture was collected and quenched by adding an equal volume of ice-cold methanol. NIC-*d3* was then added as the internal standard at a final amount corresponding to 1 μL of a 500 μM stock solution. Samples were vortexed and centrifuged, and 2 μL of the resulting supernatant was analyzed by LC-MS/MS. Nicotine-degrading activity was calculated from the decrease in nicotine concentration over time. One unit of ssNox activity was defined as the amount of catalytic activity that degraded 1 nmol of nicotine per minute at 37 °C and pH 7.4. Spore-associated activity was normalized to spore mass and expressed as units per milligram of spores.

*5.2. Catalase activity assay*

Catalase activity was measured using a Catalase Assay Kit (Beyotime Biotechnology) according to the manufacturer's instructions. Briefly, 10 μL of spore suspension was mixed with catalase assay buffer to a final volume of 40 μL. The reaction was initiated by adding 10 μL of 250 mM $H_2O_2$ and incubated at 30 °C for 15 min. The reaction was terminated by adding 450 μL of stop solution. A 10-μL aliquot of the terminated reaction mixture was diluted with 40 μL of assay buffer. Subsequently, 10 μL of the diluted sample was transferred to a 96-well plate containing 200 μL of chromogenic solution. After incubation at 30 °C for 15 min, absorbance a 570 nm was measured using a microplate reader. Catalase activity was calculated according to the kit instructions. One unit of catalase activity was defined as the amount of catalytic activity that decomposed 1 μmol of $H_2O_2$ per minute at 30 °C and pH 7.4. Spore-associated catalase activity was normalized to spore mass and expressed as units per milligram of spores.

**6. Tolerance of recombinant spores in simulated gastrointestinal fluids**

The survival and functional stability of WT and recombinant spores under simulated gastrointestinal conditions were evaluated using simulated gastric fluid containing pepsin (SGF-II) and simulated intestinal fluid containing trypsin and bile salts (SIF-II). SGF-II and SIF-II were purchased from Shanghai Yuanye Bio-Technology Co., Ltd. Spores corresponding to 3 mg dry weight were resuspended in 500 μL of SGF-II or SIF-II and incubated for the indicated durations. Proteolysis was terminated by adding 10 μL of EDTA-free protease inhibitor cocktail prepared at one tablet per 10 mL (InStab™ Protease Cocktail). Samples were centrifuged at 5,000 rpm for 5 min at room temperature, and the supernatants were discarded. The spore pellets were washed three times with sterile water to remove residual simulated fluid and were subsequently resuspended in 500 μL of PBS, pH 7.4. Residual nicotine-degrading activity was measured by adding 10 μL of 400 μM nicotine solution to the spore suspension and incubating the mixture for 20 min. A 100-μL aliquot was mixed with an equal volume of methanol to terminate the reaction and was analyzed by LC-MS/MS.

To evaluate germination capacity after treatment, a separate aliquot of each spore suspension was serially diluted and plated on selective agar. Colony counts obtained after SGF-II or SIF-II treatment were compared with those obtained from untreated spores.

**7. Growth curves of recombinant *B. subtilis***

WT and recombinant strains were inoculated into 2×YT medium and cultured overnight. The overnight cultures were diluted 1:100 into fresh 2×YT medium. $OD_{600}$ was measured every 2 h for 12 h at 0, 2, 4, 6, 8, 10, and 12 h, with sterile 2×YT medium used as the blank. Samples with $OD_{600}$ values above the linear measurement range of 0.2 to 0.8 were diluted with fresh medium before measurement. The actual $OD_{600}$ were calculated by multiplying the measured value by the corresponding dilution factor. Growth curves were generated by plotting $OD_{600}$ against incubation time.

**8. Preparation and characterization of pH-responsive spore-loaded microspheres**

pH-responsive spore-loaded microspheres, designated $rBS^{EN}$@SA&HA@CS MP, were prepared by water-in-oil emulsification and calcium -mediated cross-linking of sodium alginate and sodium hyaluronate, followed by coating with carboxymethyl chitosan. Microsphere morphology was examined by transmission electron microscopy (TEM) and scanning electron microscopy (SEM). Zeta potential was measured using a Zetasizer Ultra. pH responsiveness was evaluated by measuring the swelling ratio in buffers of different pH values. In vitro release was assessed sequentially in enzyme-free simulated gastric fluid (SGF-I), pH 2.0, and enzyme-free simulated intestinal fluid (SIF-I), pH 6.8.

*8.1. Preparation of the aqueous phase*

Sodium alginate at 3% w/v (Aladdin Biochemical Technology Co., Ltd.) and oligo-sodium hyaluronate at 1% w/v (Shandong Tiansheng Biological Technology Co., Ltd.) were dissolved in 10 mL of sterile deionized water. After complete hydration, whey protein at 3% w/v (Shanghai Yuanye Bio-Technology Co., Ltd.) and ultrafine calcium carbonate at 1% w/v were added sequentially. Purified spores were resuspended in 2% w/v mannitol, which was used as a lyoprotectant, and then added to the polymer solution. The mixture was homogenized and adjusted to a final volume of 20 mL with sterile deionized water to form the aqueous phase.

*8.2. Emulsification and calcium cross-linking*

The aqueous phase was added dropwise to 40 mL of liquid paraffin containing 1% v/v Span 80 (Aladdin Biochemical Technology Co., Ltd.). Before addition of the aqueous phase, the oil phase was stirred at 400 rpm until uniformly dispersed. The mixture was then emulsified at 600 rpm for 30 min to form a water-in-oil emulsion. The resulting emulsion was added dropwise to 40 mL of 3% w/v calcium chloride solution using a 20-mL syringe while the calcium chloride solution was continuously stirred. The microspheres were cross-linked for 1 h at room temperature and allowed to sediment. The suspension was transferred to 50-mL centrifuge tubes and centrifuged at 3,000 rpm for 5 min at 4 °C. The supernatant was discarded, and the microspheres were washed twice with sterile water.

*8.3. Carboxymethyl chitosan coating and lyophilization*

The cross-linked microspheres were resuspended in 15 mL of 0.6% w/v carboxymethyl chitosan solution, pH 6.0, with a viscosity of 265 mPa·s (Qingdao Hehai Biotechnology Co., Ltd.). The microspheres were incubated in the coating solution for 15 min. The coating procedure was performed twice, with washing between coating steps. The coated microspheres were washed and lyophilized using an Alpha 1-2 LDplus freeze-dryer (Martin Christ) for 12 to 24 h until a constant mass was obtained.

*8.4. Determination of swelling ratio (SR)*

Dried spore-loaded microspheres weighing 10 mg were immersed in 1 mL of buffer at pH 1.2, 5.5, 6.0, 6.5, 7.2, or 7.4. Samples were incubated at 37 °C for 30 min and centrifuged at 6,000 rpm for 10 min. The supernatant was removed, and excess surface liquid was gently blotted from the microspheres with filter paper before weighing. The swelling ratio was calculated as: SR=m'/m, where m' is the mass of the swollen microspheres and m is the initial dry mass. Changes in microsphere morphology at different pH values were documented by microscopy.

*8.5. In vitro release under simulated gastrointestinal conditions*

Spore-loaded microspheres weighing 10 mg were dispersed in 1 mL of SGF-I and incubated at 37 °C with agitation at 100 rpm. At 1 and 2 h, 100-μL samples of the supernatant were collected, serially diluted 10-fold to $10^{-5}$, and plated on selective agar for colony enumeration. After 2 h in SGF-I, the samples were centrifuged at 6,000 rpm for 10 min. The supernatant was removed, and the microsphere pellet was transferred to 1 mL of SIF-I. Incubation was continued at 37 °C and 100 rpm. Supernatant samples were collected at 2, 3, 4, and 5 h after transfer to SIF-I, serially diluted, and plated as described above. An equivalent amount of free spores was subjected to the same sequential SGF-I and SIF-I treatments as a control. Released spores were quantified by colony counting, and cumulative release was calculated relative to the total viable spore content of the initial formulation. Unencapsulated spores remaining in the supernatant after microsphere preparation were also quantified to determine encapsulation efficiency.

**9. Biochemical analysis of serum and tissue homogenates**

Blood samples were obtained by retro-orbital bleeding. For serum preparation, whole blood was collected into microcentrifuge tubes without anticoagulant and allowed to clot on ice or at 4 °C for 2 h. Samples were then centrifuged at 3000 rpm for 5 min. The supernatant was carefully transferred to a new tube and centrifuged again under the same conditions to remove residual cells and debris. Hemolyzed samples were excluded from subsequent analyses. Serum aliquots were stored at −20 or −80 °C until use. For plasma preparation, whole blood was collected directly into tubes containing sodium heparin as the anticoagulant. Before LC–MS/MS analysis, serum or plasma samples were mixed with three to four volumes of methanol to precipitate proteins. After vortexing and centrifugation, the clear supernatant was collected for quantitative LC–MS/MS analysis.

For tissue homogenate preparation, tissues were rinsed with ice-cold PBS (pH 7.4) to remove residual blood, weighed, and minced. Tissue samples were homogenized in PBS at a ratio of 100 mg tissue per 1 mL PBS using a low-temperature tissue grinder (Shanghai Jingxin) with three to four ceramic beads. The homogenates were first centrifuged at 1,000 rpm for 10 min, and the resulting supernatants were transferred to new tubes and centrifuged at 12,000 rpm for 15 min. The final supernatants were collected for biochemical analyses. Whole-brain tissue was used to prepare brain homogenates. For RNA extraction, tissue samples were homogenized directly in 1 mL Trizol reagent (Accurate Biology).

Total bilirubin (TBIL) was measured using a colorimetric assay kit (Elabscience). Aspartate aminotransferase (AST), alanine aminotransferase (ALT), creatinine (Cre), and blood urea nitrogen (BUN) were measured using commercial assay kits from Nanjing Jiancheng Bioengineering Institute. IL-6, TGF-β1, and dopamine (DA) concentrations were determined using ELISA kits from Boyan Biology.

**10. Evaluation of the nicotine-degrading activity of spore microspheres**

To evaluate the nicotine-degrading activity, 15-mg of lyophilized spore microspheres was weighed and allowed to swell in 300 μL PBS for 10 min. The reaction was initiated by adding 200 μL nicotine solution at a concentration of 0.05 mg/mL. Reaction were conducted under physiological conditions at 37 °C and pH 7.4. At 0, 5, 10, 20, 30, 40, 50, and 60 min, 100-μL samples were collected. Each sample was immediately mixed with two volumes of methanol to terminate the reaction, followed by the addition of 1 μL deuterated nicotine internal standard (NIC-*d3*, 500 μM). Samples were vortexed thoroughly and centrifuged, and 2 μL of the resulting supernatant was injected into the LC–MS/MS system for nicotine quantification. The sample collected immediately after reaction initiation at 0 min was used as the baseline control. One activity unit was defined as the amount of catalytic activity in 10 mg of spore microspheres that degraded 1 nmol nicotine per minute at 37 °C and pH 7.4.

#### 11. Neurobehavioral assessments

*11.1. Optimization of the oral nicotine dose*

To minimize potential differences in consumption or responses caused by the bitter taste of nicotine, all nicotine solutions were prepared in 0.1% sodium saccharin. Before behavioral testing, mice were habituated to the saccharin solution for 3 d to reduce novelty-related stress. C57BL/6 mice were randomly assigned to receive vehicle (0.1% sodium saccharin) or nicotine at 0.5, 1.0, or 1.5 mg/kg by oral gavage (n = 3 per group). Fifteen minutes after administration, each mouse was placed in an empty chamber for acclimation and subsequently positioned in the center of an open-field arena. Locomotor activity was recorded for 10 min using the ANY-maze video-tracking system (Stoelting Co., USA) to determine the nicotine dose that induced acute locomotor hyperactivity. The arena was cleaned with 75% ethanol between trials to remove residual olfactory cues.

*11.2. Pharmacodynamic evaluation of spore microspheres following acute nicotine administration*

Mice were randomly assigned to the indicated treatment groups (n = 8 per group). To maintain uniform dispersion and minimize sedimentation, $rBS^{EN}$@SA&HA@CS microspheres were resuspended in 0.5% sodium carboxymethyl cellulose (CMC-Na) immediately before administration. Control mice received equivalent volumes of CMC-Na and sodium saccharin solutions. Mice in the treatment group received $rBS^{EN}$@SA&HA@CS microspheres (15 mg) by oral gavage 1 h before nicotine administration. This interval was selected to allow the microspheres to reach the proximal intestine and initiate the release of the engineered spores. Freshly prepared nicotine was then administered by oral gavage at 0.5 mg/kg. Fifteen minutes after nicotine administration, mice were subjected to the open-field test. Behavior was recorded for 5 min using a Gene&I video-tracking system and analyzed using DigBehv software. Total distance traveled, mean velocity, and time spent in the center zone were quantified to evaluate the effects of the spore microspheres on nicotine-induced behavioral alterations.

#### 12. Establishment of the nicotine addiction model for evaluations

*12.1. Nicotine addiction model*

The nicotine addiction model was established by repeated subcutaneous (s.c.) administration of nicotine using protocols adapted from Akinola and Kim et al. (10, 11). Nicotine was dissolved in sterile physiological saline, and fresh solutions were prepared immediately before administration. Varenicline, an approved smoking-cessation medication, was used as a positive control. C57BL/6 mice were randomly assigned to five groups (n = 6 per group): (a) vehicle control, (b) NIC model, (c) rBS$^{EN}$@SA&HA@CS MP + NIC preventive treatment, (d) NIC + rBS$^{EN}$@SA&HA@CS MP therapeutic treatment, and (e) NIC + varenicline. During the initial sensitization phase (Days 1–9), mice in the NIC model, therapeutic-treatment, and varenicline groups received nicotine once daily by s.c. injection at 0.5 mg/kg. Mice in the preventive-treatment group received rBS$^{EN}$@SA&HA@CS MP (15 mg) by oral gavage 1 h before each daily nicotine injection. Vehicle-control mice received corresponding injections of saline and oral administration of the blank vehicle. On day 10, all treatments were withheld for 24 h to induce spontaneous nicotine withdrawal, after which the mechanical paw withdrawal threshold (PWT) was measured. During the subsequent dose-escalation phase (Days 11–13), the nicotine dose was increased to 1.0 mg/kg in all nicotine-treated groups. At the same time, mice in the therapeutic-treatment group began receiving rBS$^{EN}$@SA&HA@CS MP (15 mg) once daily by oral gavage, whereas mice in the varenicline group received varenicline at 0.5 mg/kg by oral gavage once daily. On day 14, all treatments were discontinued, and behavioral assessments and final PWT measurements were performed to evaluate treatment efficacy.

*12.2. Nicotine withdrawal*

Nicotine withdrawal can be assessed using either spontaneous or pharmacologically precipitated paradigms. Mecamylamine (MEC; 2 mg/kg) is commonly used to precipitate nicotine withdrawal. However, MEC has been reported to exert antidepressant-like effects (11), which may confound the evaluation of withdrawal-associated depressive behaviors. In preliminary experiments using nicotine-naive mice, intraperitoneal administration of MEC produced marked sedation and reduced locomotor activity within 10 min (Fig. S24A), indicating a direct suppressive effect on the central nervous system. MEC also increased the baseline PWT (Fig. S24B), suggesting an analgesic-like effect that could mask withdrawal-induced mechanical hypersensitivity and thereby produce false-negative results. We therefore used spontaneous withdrawal rather than MEC-precipitated withdrawal in subsequent experiments.

*12.3. Measurement of the mechanical paw withdrawal threshold (PWT) using the von Frey filaments*

Nicotine withdrawal is accompanied by somatic signs, affective disturbances, and increased pain sensitivity. Withdrawal-associated mechanical hypersensitivity was quantified by measuring the hind-paw withdrawal threshold using the von Frey filaments and an Aesthesio® Precision Tactile Sensory Evaluator. Mice were individually placed in transparent Plexiglas chambers positioned on an elevated wire-mesh platform with a 1 × 1-cm grid. Testing was conducted in a quiet, temperature-controlled room under dim lighting. Mice were allowed to acclimate for 20 min or until exploratory behavior had subsided. Mechanical sensitivity was assessed using the up-down method. Testing was initiated with a filament of intermediate bending force. Each filament was applied perpendicularly to the midplantar surface of the hind paw from beneath the mesh, avoiding the footpads and digits. The filament was advanced until it buckled and was maintained in position for 2–4 s. The stimulation site was varied between trials to minimize local sensitization or desensitization, and stimuli were delivered at irregular intervals of at least 10 s. A brisk paw withdrawal, paw lifting, or licking of the stimulated area was scored as a positive response. An absence of movement or a minor postural adjustment was scored as a negative response. Following a negative response, the next filament of greater force was applied; following a positive response, the next filament of lower force was used. Each selected filament was applied five times. The corresponding force was defined as the PWT when at least three positive responses were observed in five applications. If all five applications produced negative responses, mice were allowed to rest for 1 min before testing with the next filament of greater force. All measurements were performed by an investigator blinded to the experimental group assignments.

#### 13. *In vivo* and *ex vivo* near-infrared fluorescence imaging

##### *13.1. Fluorescence labeling and validation*

A 15-mg aliquot of the spore microspheres was resuspended in 500 μL of deionized water and incubated with DiR dye (Yeasen Biotechnology) at a final concentration of 10 μM. The suspension was incubated at 37 °C for 1 h with shaking and protected from light. Labeled microspheres were collected by centrifugation at 5,000 rpm for 5 min and washed three times with deionized water to remove unbound dye. DiR labeling was confirmed by fluorescence microscopy using an IX73 inverted fluorescence microscope (Olympus) after the microspheres were transferred to a six-well plate.

##### *13.2. In vivo spatiotemporal tracking and biodistribution*

To reduce interference from abdominal fur during fluorescence imaging, the abdominal region of each C57BL/6 mouse was shaved and treated with depilatory cream 24 h before imaging. Mice were administered 15 mg of DiR-labeled microspheres by oral gavage and were allowed free access to water thereafter. Whole-body fluorescence imaging was performed at 1, 3, 6, 12, and 24 h after administration using an IVIS Lumina III imaging system (PerkinElmer). Mice were anesthetized with isoflurane in an induction chamber and maintained under isoflurane anesthesia during imaging. Mice administered unlabeled microspheres served as negative controls (n = 3 per group). After completion of the final in vivo imaging session, mice were euthanized by cervical dislocation. The heart (H), liver (L), spleen (S), lung (P), kidney (K), brain

(B), large intestine (LI), and small intestine (SI) were collected and imaged ex vivo to assess the tissue distribution of the fluorescence signal.

**14. Cellular protein extraction and quantification**

After removal of the culture medium, cells were washed twice with ice-cold PBS. Ice-cold cell lysis buffer (60–80 μL; Beyotime Biotechnology, Shanghai, China) was then added to each culture dish. The cells were detached using a cell scraper, transferred to microcentrifuge tubes, and incubated on ice for an additional 10 min to ensure complete lysis. The lysates were centrifuged at 12000 rpm for 10 min at 4 °C, and the supernatants containing total cellular proteins were transferred to pre-chilled microcentrifuge tubes. Protein concentrations were determined using a BCA Protein Assay Kit (Generay Biotech, Shanghai, China) according to the manufacturer's instructions. Protein lysates from the different treatment groups were subsequently adjusted to the same concentration before downstream analyses. Subsequent procedures were performed as previously described (12).

**15. Histological sectioning and hematoxylin and eosin staining**

Mice were anesthetized by intraperitoneal injection of 1% sodium pentobarbital, and blood was collected from the retro-orbital sinus. The mice were subsequently euthanized by cervical dislocation. The heart, liver, spleen, lung, kidney, brain, and intestines were collected and gently rinsed with PBS to remove residual blood. The intestinal lumen was additionally flushed with PBS to remove luminal contents. The tissues were completely immersed in a 4% paraformaldehyde (Servicebio, Wuhan, China) and processed for paraffin embedding, sectioning, and hematoxylin and eosin (H&E) staining by Jiangsu Ketu Biotechnology Co., Ltd. After ethanol dehydration and paraffin embedding, the tissues were sectioned at a thickness of 4 μm. The sections were deparaffinized by sequential immersion in xylene twice for 15 min each, anhydrous ethanol for 5 min, 85% ethanol for 5 min, and 75% ethanol for 5 min, followed by washing with tap water. The sections were stained with hematoxylin for 4 min, rinsed with tap water to remove excess stain, briefly differentiated in differentiation solution, and treated with bluing solution. The sections were then immersed in 85% ethanol for 4 min and 95% ethanol for 4 min, followed by eosin staining for 3 min. After staining, the sections were dehydrated and cleared by sequential immersion in anhydrous ethanol three times for 4 min each, n-butanol twice for 4 min each, and xylene twice for 5 min each. The sections were mounted with neutral balsam and covered with coverslips. Whole-slide images were acquired using a Pannoramic slide scanner and analyzed using CaseViewer 2.4 software. Nuclei were stained blue, whereas the cytoplasm and extracellular components were stained various shades of pink to red.

#### 16. 16S rRNA sequencing and gut microbiota analysis

##### *16.1. Sample collection and processing*

Fresh fecal pellets were collected at the designated experimental time points directly into sterile tubes, with three to four pellets obtained from each mouse. Samples were immediately frozen in liquid nitrogen and stored at -80 °C until analysis. Genomic DNA extraction, library preparation, and high-throughput sequencing were performed by LC-Bio Technology Co., Ltd. (Hangzhou, China). The libraries were sequenced on an Illumina NovaSeq 6000 platform. Data visualization and selected downstream analyses were performed using the OmicStudio cloud platform.

##### *16.2. DNA extraction and PCR amplification*

Total genomic DNA was extracted from the fecal samples using a fecal genomic DNA extraction kit according to the manufacturer's instructions. DNA concentration was measured using a Qubit fluorometer, and DNA quality was assessed before library preparation. The V3–V4 hypervariable region of the bacterial 16S rRNA gene was amplified using the universal primers 341F (5'-CCTACGGGNGGCWGCAG-3') and 805R (5'-GACTACHVGGGTATCTAATCC-3'). PCR amplification was performed with an initial denaturation at 98 °C for 30 s, followed by 32 cycles of denaturation at 98 °C for 10 s, annealing at 54 °C for 30 s, and extension at 72 °C for 45 s, with a final extension at 10 min. PCR products were purified using AMPure XT magnetic beads and quantified. Library integrity and concentration were evaluated using an Agilent 2100 Bioanalyzer and an Illumina library quantification kit. Qualified libraries were pooled at equimolar concentrations and subjected to paired-end sequencing.

##### *16.3. Sequencing data processing and bioinformatics analysis*

Raw de-multiplexed sequence reads were processed using cutadapt v1.9 to remove primer sequences. Paired-end reads were merged using FLASH v1.2.8. Quality filtering was performed using fqtrim v0.94 with a sliding-window approach. Reads with a quality score below 20, a length below 100 bp, or more than 5% ambiguous bases were removed. Chimeric sequences were identified and excluded using VSEARCH v2.3.4. The remaining sequences were denoised using DADA2 to generate amplicon sequence variants (ASVs). Representative ASV sequences were taxonomically assigned against the SILVA and NT-16S reference databases using the feature-classifier plugin in QIIME 2. Alpha- and beta-diversity metrics were calculated in QIIME 2, and microbial community composition was evaluated based on taxonomic relative abundance. Differences in genus-level relative abundance between groups were assessed using the

Wilcoxon rank-sum test, with $P < 0.05$ considered statistically significant. Linear discriminant analysis effect size (LEfSe) was used to identify differentially abundant taxa, with an LDA score of at least 3.0 and $P < 0.05$. Graphical analyses were performed using the appropriate packages in R v3.4.4.

**17. Quantitative real-time PCR analysis of nAChR subunit expression**

The mRNA expression levels of the $\alpha$4, $\beta$2, and $\alpha$7 nAChR subunits were measured by quantitative real-time PCR (qPCR), with GAPDH used as the reference gene. Total RNA was extracted from the indicated tissue samples. One microgram of total RNA was subjected to genomic DNA removal and first-strand cDNA synthesis using a reverse-transcription kit from Accurate Biology (Hunan, China) according to the manufacturer's instructions. Quantitative PCR was performed using a qPCR master mix from the same manufacturer on a LightCycler® 96 Real-Time PCR System (Roche). Fluorescence data were collected and analyzed using the accompanying instrument software. The primer sequences were as follows: $\alpha$4 (forward: 5'-CTAGCAGCCACATAGAGAC-3', reverse: 5'-GACAAGCCAAAGCGGACAAG-3'), $\beta$2 (forward: 5'-GAGGTGAAGCACTTCCCATTT-3', reverse: 5'-GCCACATCGCTTTTGAGCAC-3'), $\alpha$7 (forward: 5'-CCGTGCCCTTGATAGCACA-3', reverse: 5'-GGCATTTTGCCACCATCAGG-3'), and GAPDH (forward: 5'-AGGTCGGTGTGAACGGATTTG-3', reverse: 5'-TGTAGACCATGTAGTTGAGGTCA-3').

**18. Nissl staining and cleaved caspase-3 immunofluorescence of brain tissue sections**

Brain sections were subjected to Nissl staining to assess neuronal morphology and tissue architecture. Sections were stained using a commercial Nissl staining solution (Beyotime) according to the manufacturer's instructions and examined for changes in neuronal morphology and Nissl body abundance. For cleaved caspase-3 immunofluorescence, adjacent brain sections were deparaffinized and rehydrated through a graded ethanol series. Antigen retrieval was performed by heating the sections in sodium citrate buffer. After blocking nonspecific binding, the sections were incubated overnight at 4 °C with a primary antibody against cleaved caspase-3. Following washing, the sections were incubated with the appropriate secondary antibody for 1 h at room temperature. Nuclei were counterstained with DAPI. Images were acquired using a fluorescence microscope and analyzed to evaluate cleaved caspase-3 immunoreactivity and associated histopathological changes.

### Supporting Figures

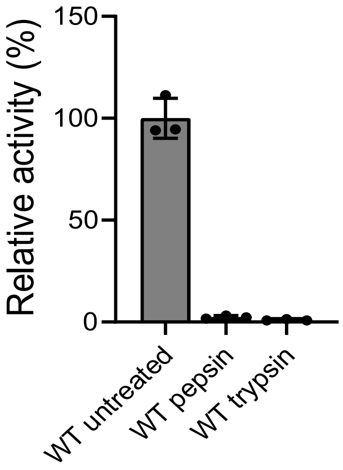


**Figure S1.** Relative activity of WT ssNox after the protease treatments. Data are presented as mean ± SD (n = 3).

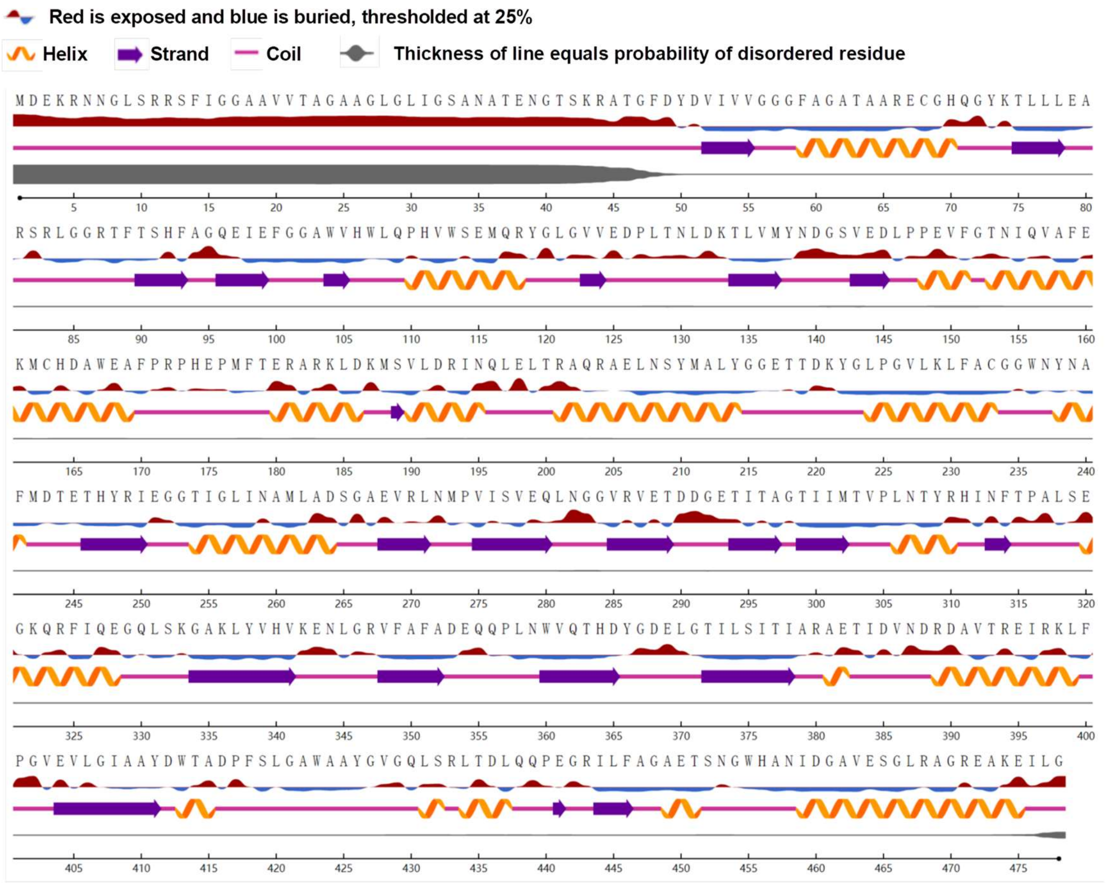


**Figure S2.** Secondary structure and relative solvent accessibility of ssNox obtained by using NetSurfP-3.0.

#### Site-directed mutagenesis

Mutations: Single Point

#### Stability of designed mutations in kcal/mol

Stabilizing mutations Destabilizing mutations
Energy is in kcal/mol

| Chain | Position | Residue | Ala | Arg | Asn | Asp | Cys | Gln | Glu | Gly | His | Ile | Lys | Leu | Met | Phe | Pro | Ser | Thr | Val | Trp | Tyr |
|---|---|---|---|---|---|---|---|---|---|---|---|---|---|---|---|---|---|---|---|---|---|---|
| A | 322 | Lys | 1.1 | - | - | - | - | - | - | - | 2.2 | - | - | - | - | - | - | - | - | -0.1 | - | - |
| A | 424 | Ala | - | - | - | - | 7.4 | - | - | - | - | - | - | 184.1 | 13.0 | - | 16.7 | - | - | 45.9 | - | - |
| A | 456 | His | 3.0 | 15.3 | - | - | - | - | - | 4.9 | - | - | - | - | - | - | - | 3.4 | - | 2.8 | - | - |
| A | 330 | Gln | - | - | 0.1 | - | - | - | - | - | 6.0 | -0.1 | - | - | - | - | - | - | -0.6 | 0.4 | - | - |
| A | 311 | His | - | 2.1 | - | 2.6 | - | - | - | 2.5 | - | - | - | - | - | - | - | 2.2 | 1.6 | - | - | - |

**Figure S3.** Predicted effects of designed mutations on ssNox stability, as evaluated by HotSpot Wizard 3.0.

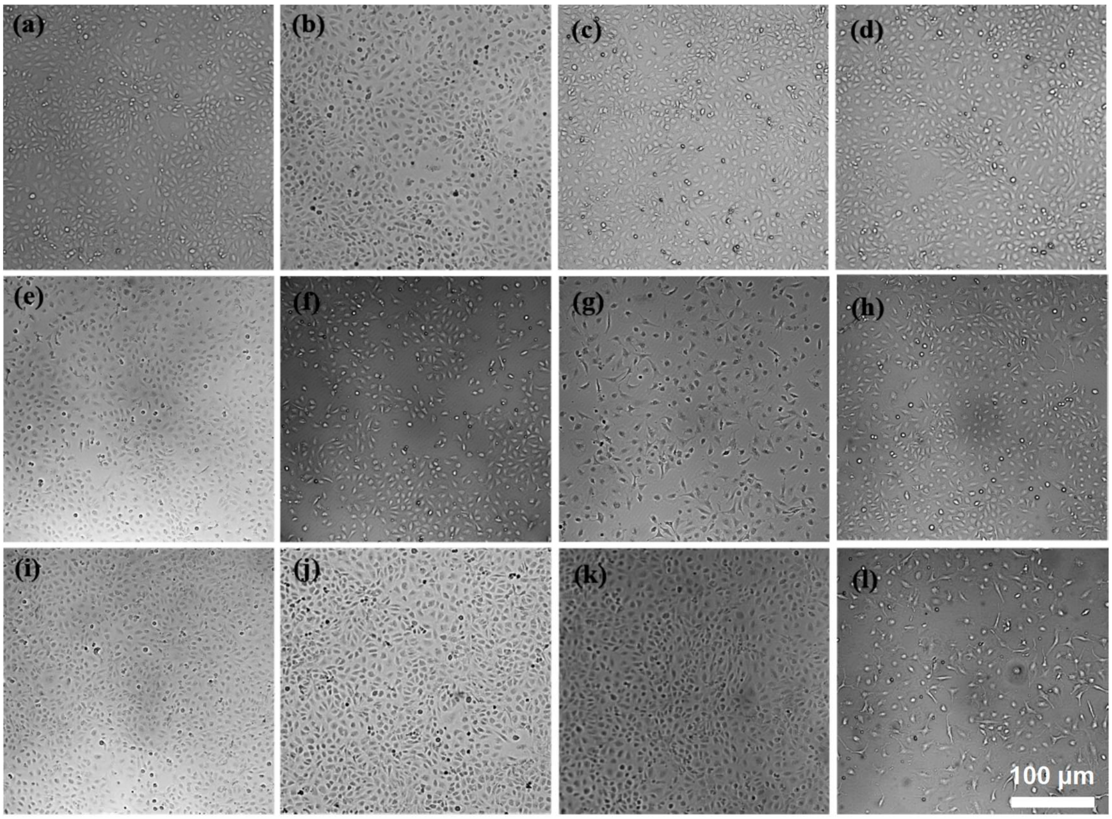


**Figure S4.** Morphology of IEC-6 cells following various treatments. The treatment conditions in panels (a)–(l) correspond to those detailed in Fig. 1H and 1I.

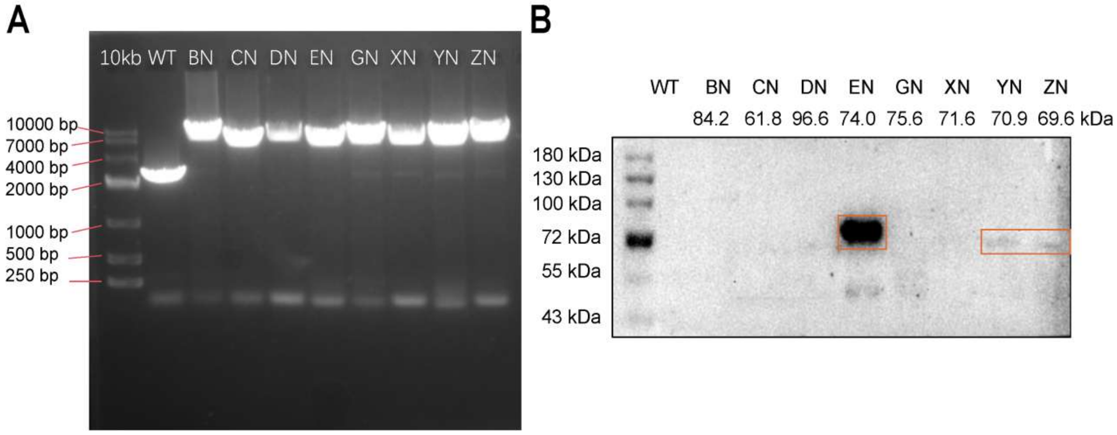


**Figure S5.** Genotypic and protein-level validation of the recombinant *B. subtilis* strains. (A) PCR verification of genomic integration in the WT and recombinant strains. (B) Corresponding Western blot analysis of the spore-displayed proteins. Strain abbreviations are the same as those described in Fig. 2B and 2C.

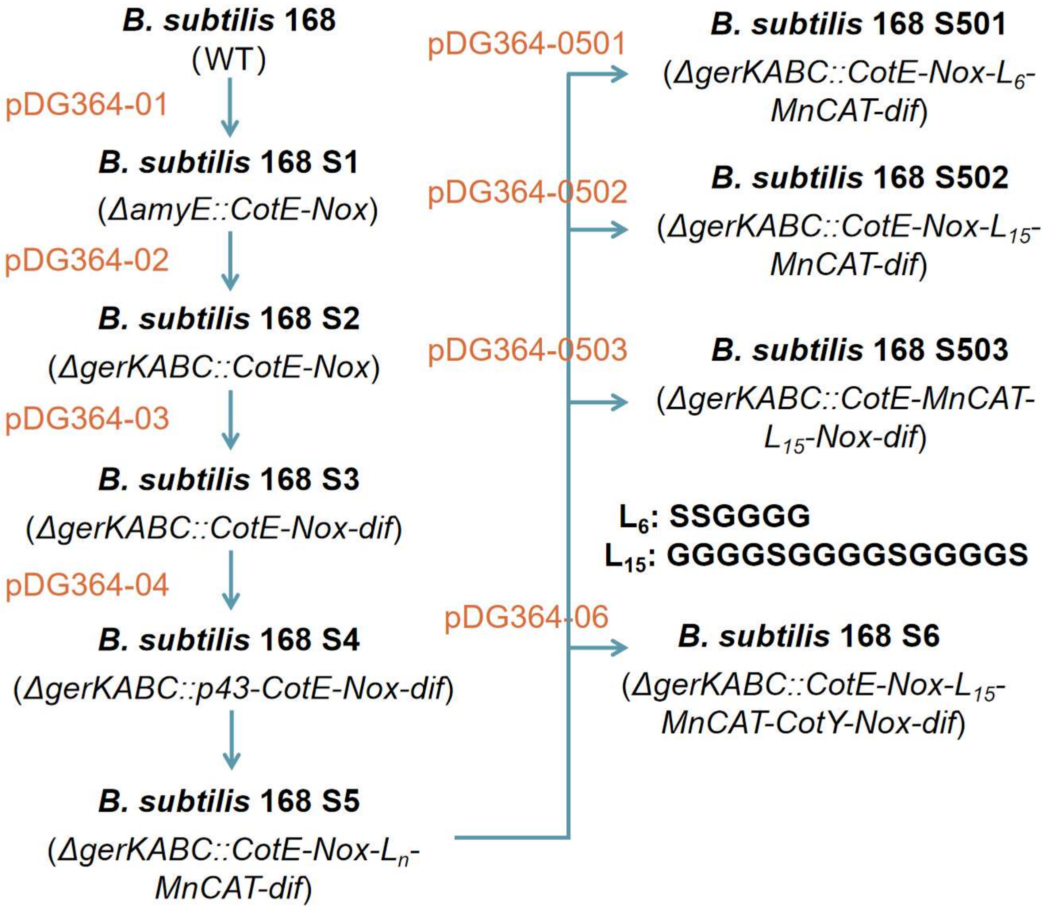


**Figure S6.** Construction workflow and the lineage of recombinant spore-forming strains used in this study. See Table S2 for details.

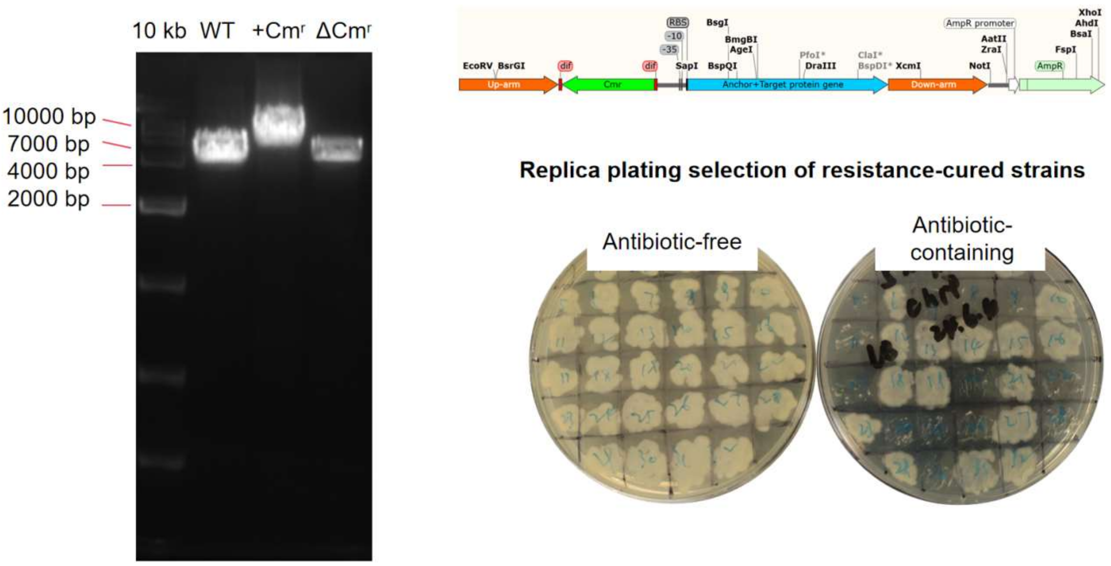


**Figure S7.** Genotypic and phenotypic validation of antibiotic-resistance marker removal. Strains retaining the resistance cassette are denoted as +Cmr, while those in which the cassette has been excised are denoted as ΔCmr.

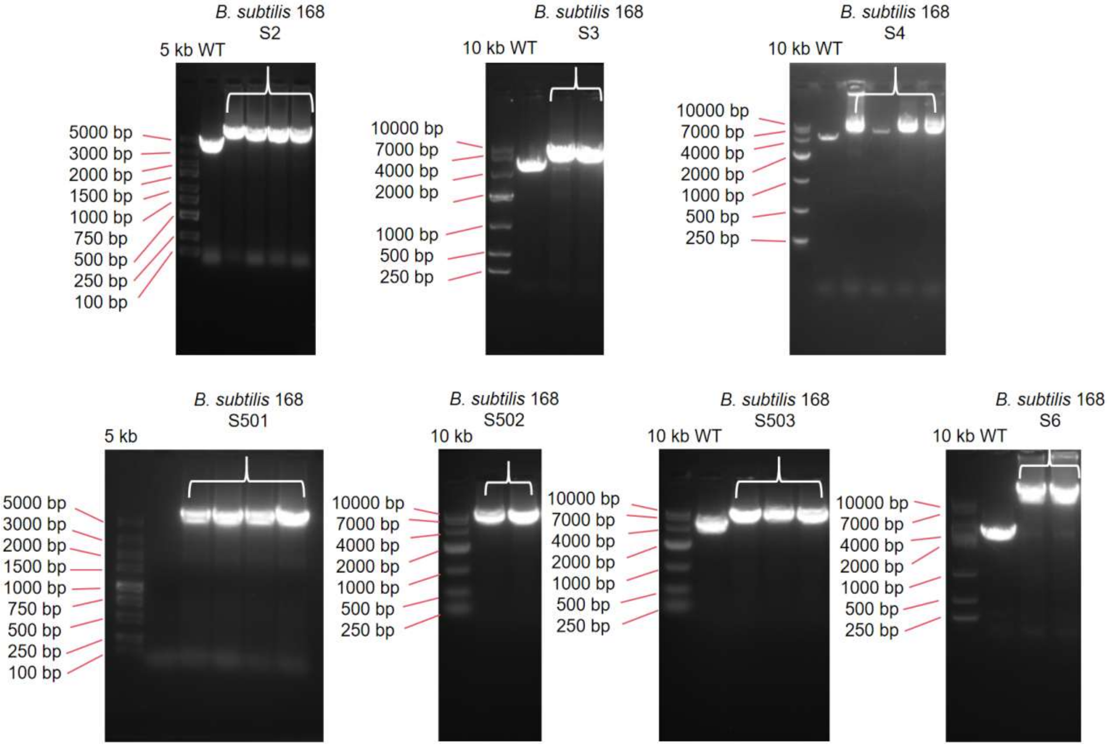


**Figure S8.** Genomic PCR verification of engineered *B. subtilis* strains S2, S3, S4, S501, S502, S503 and S6.

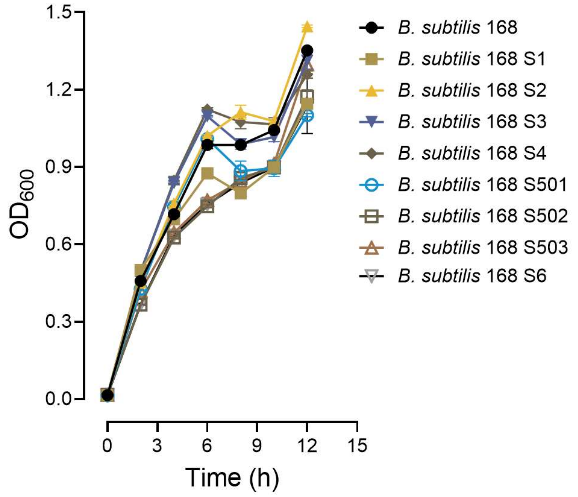


**Figure S9.** Comparative growth curves of various recombinant *B. subtilis* strains.

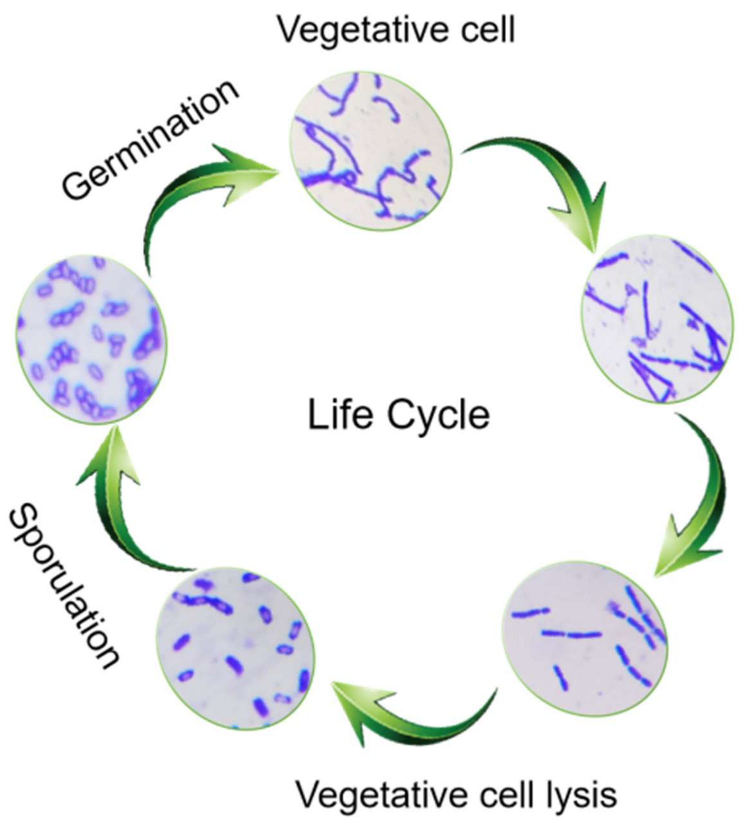


**Figure S10.** Schematic representation of *B. subtilis* life cycle.

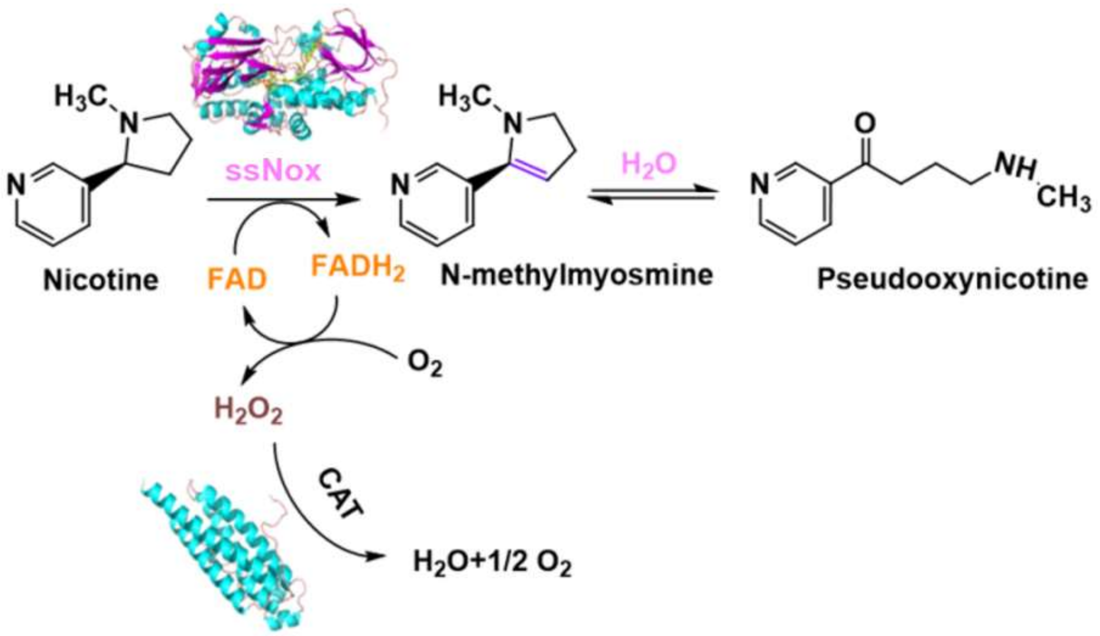


**Figure S11.** Coupled enzymatic reactions catalyzed by ssNox and CAT.

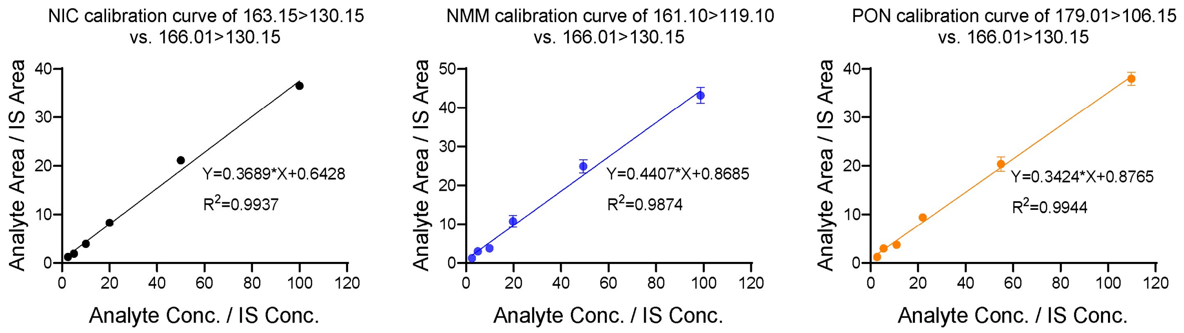


**Figure S12.** Standard calibration curves for nicotine (NIC), *N*-methylmyosmine (NMM) and pseudooxynicotine (PON) used for quantitative analysis by LC/MS/MS.

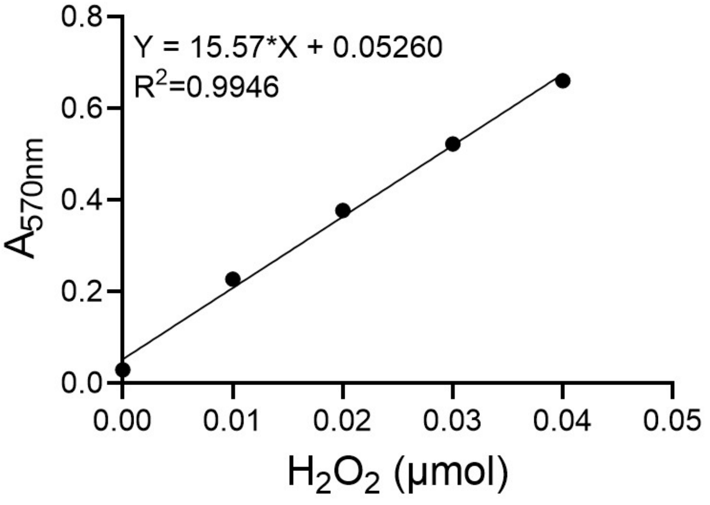


**Figure S13.** Standard calibration curve for $H_2O_2$ quantification.

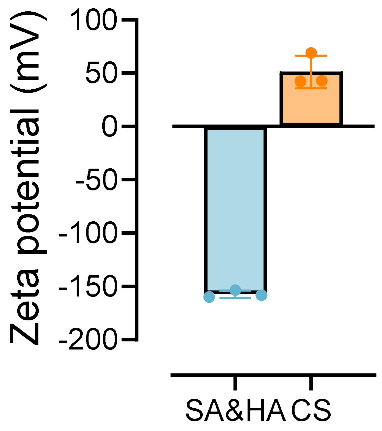


**Figure S14.** Zeta potential of the SA&HA mixture and CS solution.

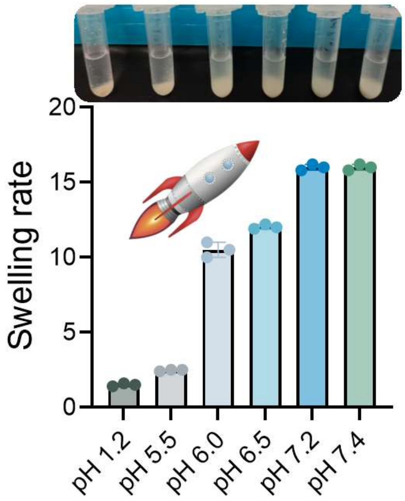


**Figure S15.** pH-dependent swelling behaviors of the microspheres.

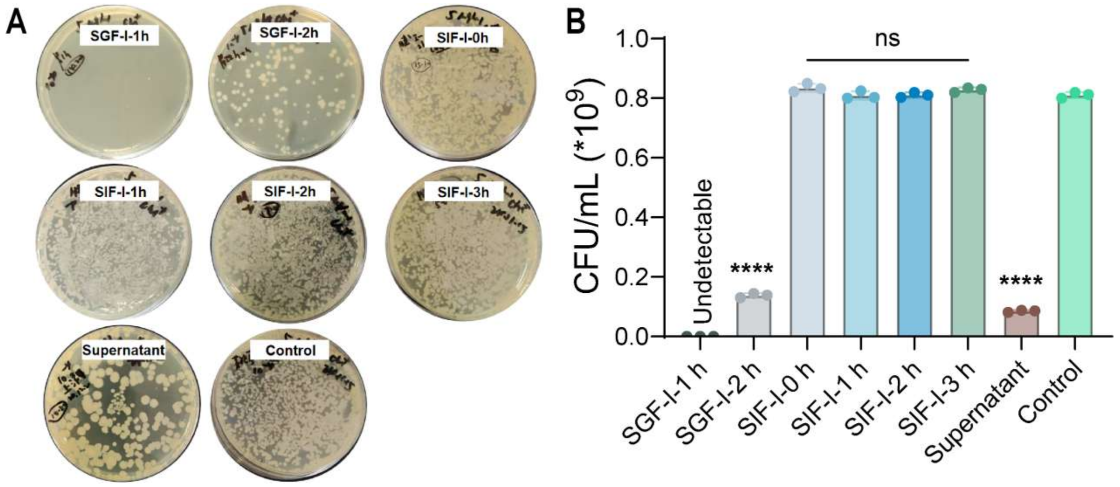


**Figure S16.** *In vitro* release profiles of the microspheres in SGF-I and SIF-I. (A) Representative images of the dilution plates used to evaluate spore release. (B) Corresponding quantitative statistical analysis evaluated by ordinary one-way ANOVA. "Supernatant" denotes the residual liquid collected immediately after microsphere preparation, whereas "Control" denotes an equivalent number of unencapsulated spores. SIF-I-0 h, SIF-I-1 h, SIF-I-2 h, and SIF-I-3 h indicate the duration of incubation after the microspheres were transferred from SGF-I to SIF-I. * $p < 0.05$, ** $p < 0.01$, **** $p < 0.0001$. Data were analyzed by ordinary one-way ANOVA (n = 3).

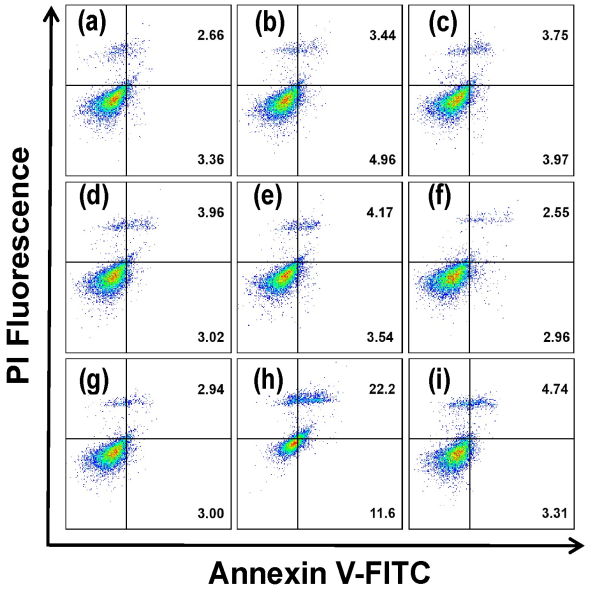


**Figure S17.** Representative flow cytometry plots showing apoptosis following co-treatment with microspheres and nicotine. Panels (a)–(i) correspond to the experimental groups detailed in Fig. 3H.

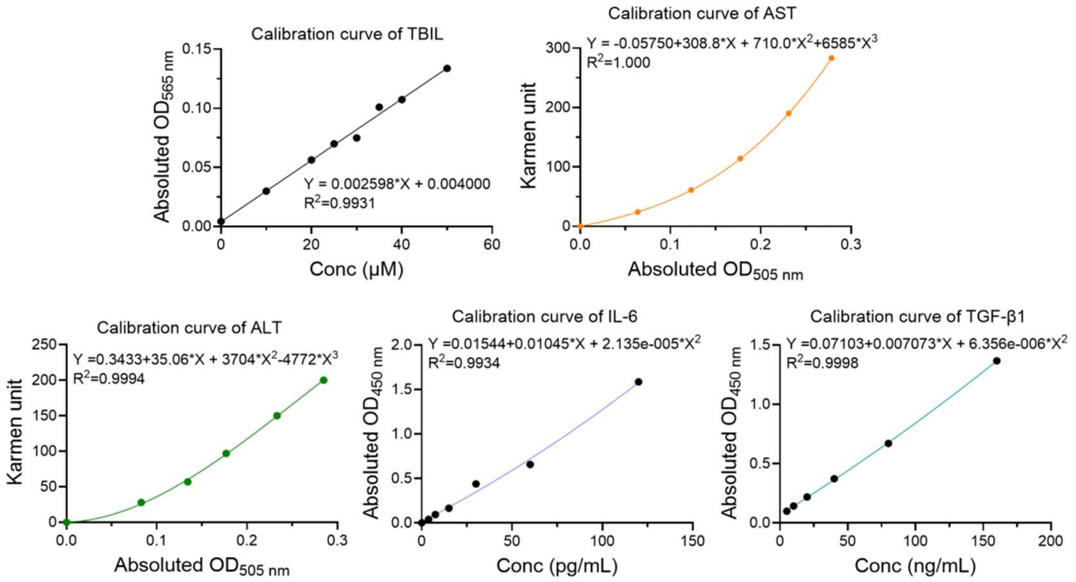


**Figure S18.** Standard calibration curves for TBIL, AST, ALT, IL-6 and TGF-β1 quantification.

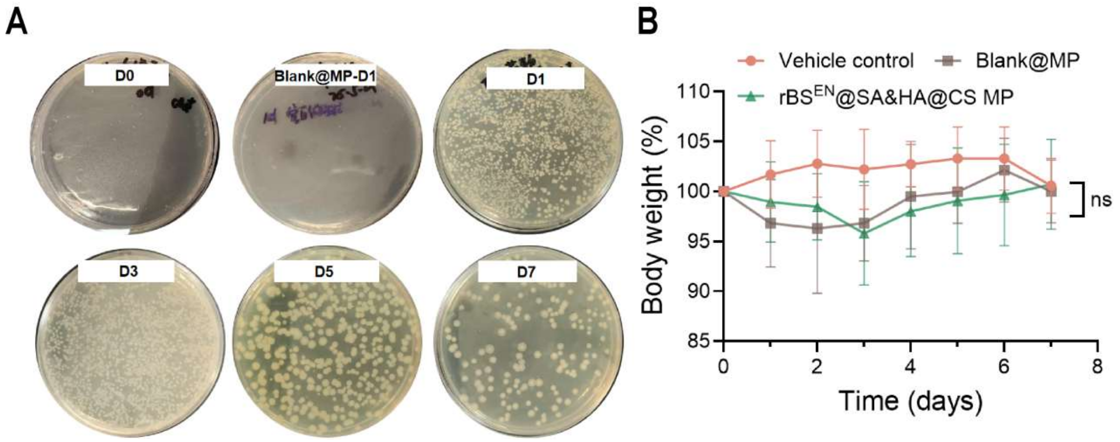


**Figure S19.** Fecal bacterial recovery and body weight during gut microbiota study. (A) Representative images of fecal dilution plates used bacterial enumeration. (B) Body weight of mice during the study (n = 8 per group). ns, not significant.

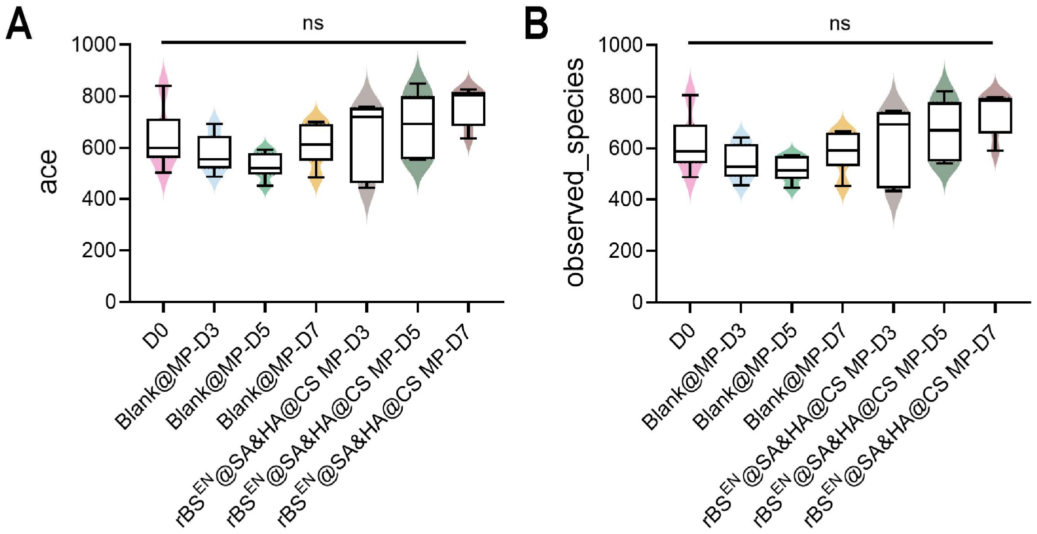


**Figure S20.** Alpha diversity indices across treatment groups and different time points. (A) Ace index. (B) Observed species index. Data were obtained from mice per treatment group (n = 6). ns, not significant.

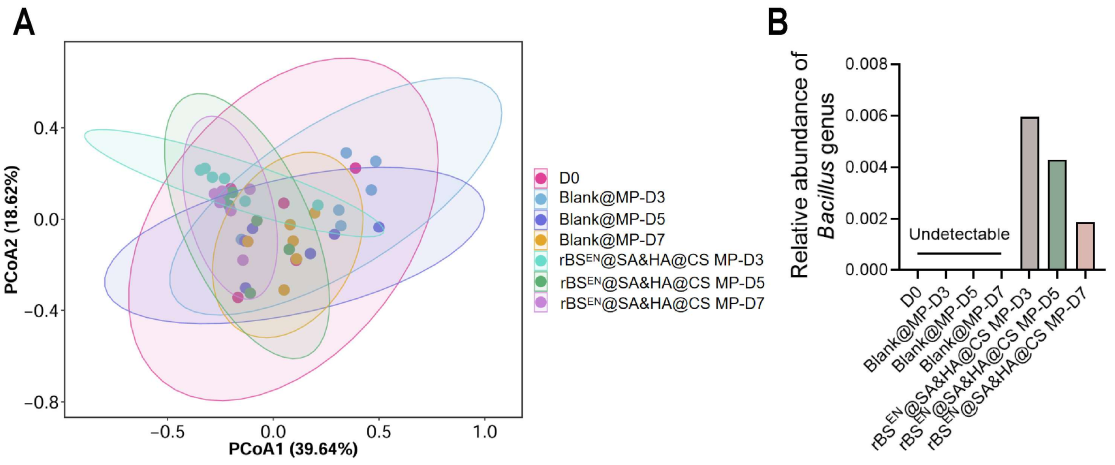


**Figure S21.** Beta diversity and taxonomic composition of the gut microbiota. (A) Principal coordinates analysis (PCoA) of microbial community composition across treatment groups and different time points. (B) Relative abundance of *Bacillus* determined by 16S rRNA sequencing.

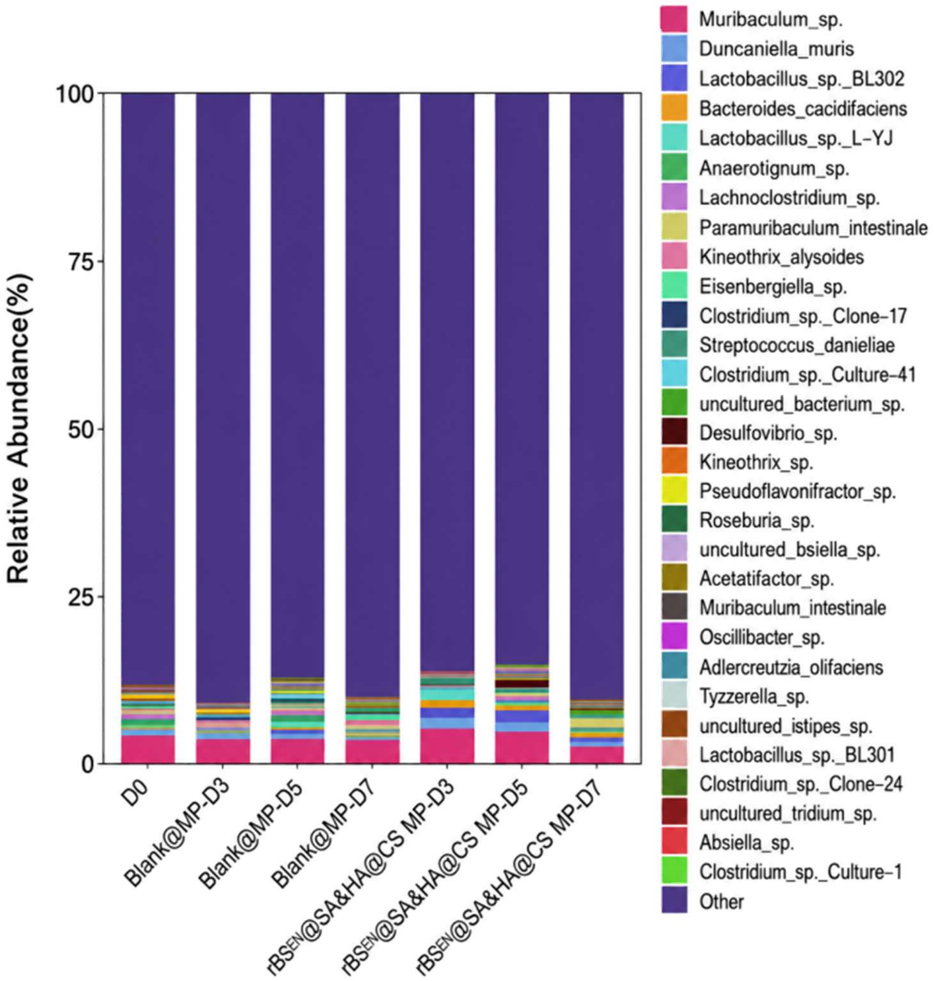


**Figure S22.** Relative abundance of top 30 microbial species across treatment groups and different time points.

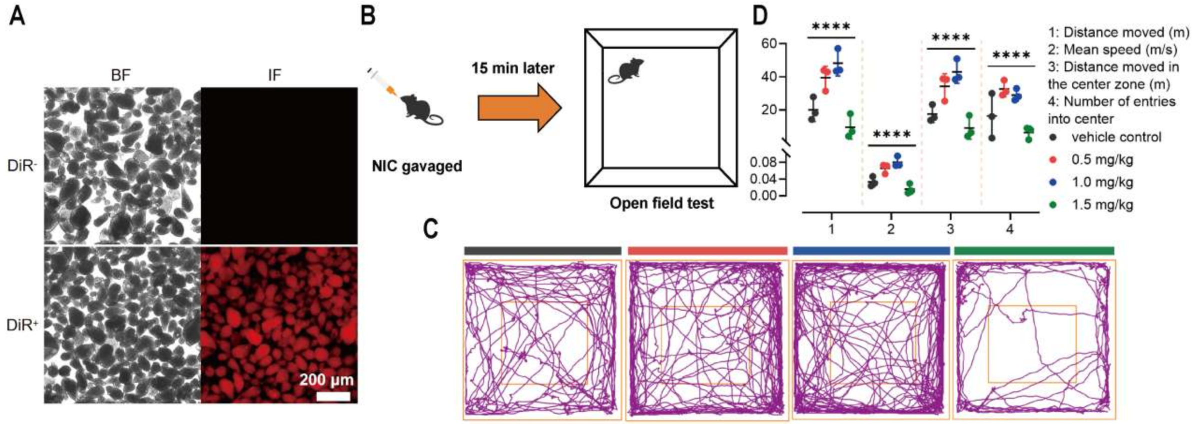


**Figure S23.** DiR-labeled microspheres and assessment of locomotor activity in the open-field test. (A) Fluorescence microscopic images of DiR-labeled microspheres. (B) Schematic illustration of the OFT. (C) Representative movement trajectories of mice in each group. (D) Quantitative analysis of locomotor activity, including total distance traveled, mean speed, distance traveled in the center zone, and number of center-zone entries (n = 3). * $p < 0.05$, ** $p < 0.01$, **** $p < 0.0001$.

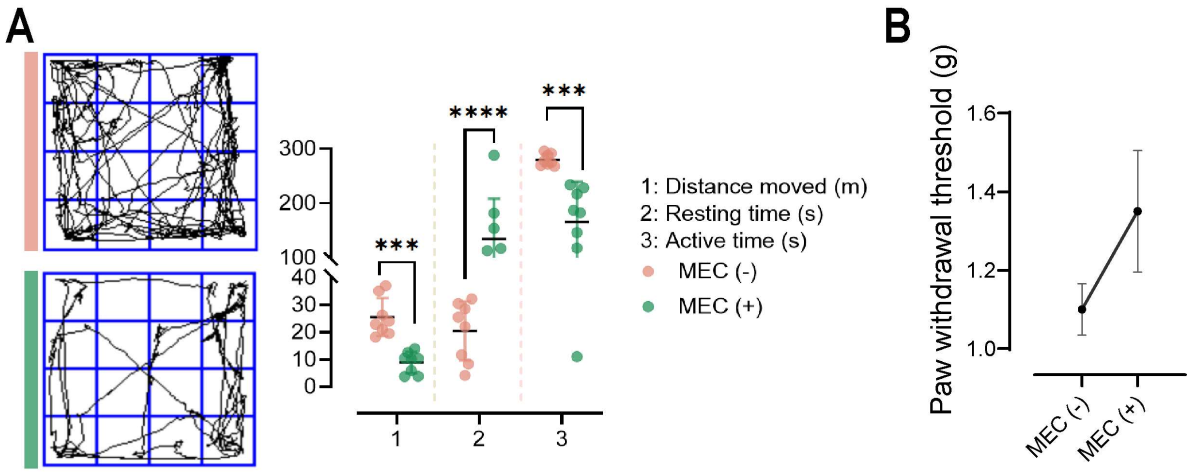


**Figure S24.** Behavioral and sensory responses following mecamylamine (MEC) treatment. (A) OFT results for naive mice 10 min after intraperitoneal administration of MEC. Data are presented as mean ± SD (n = 8) and were analyzed using two-way ANOVA. * $p < 0.05$, ** $p < 0.01$, **** $p < 0.0001$. (B) Comparison of PWTs before and after MEC administration.

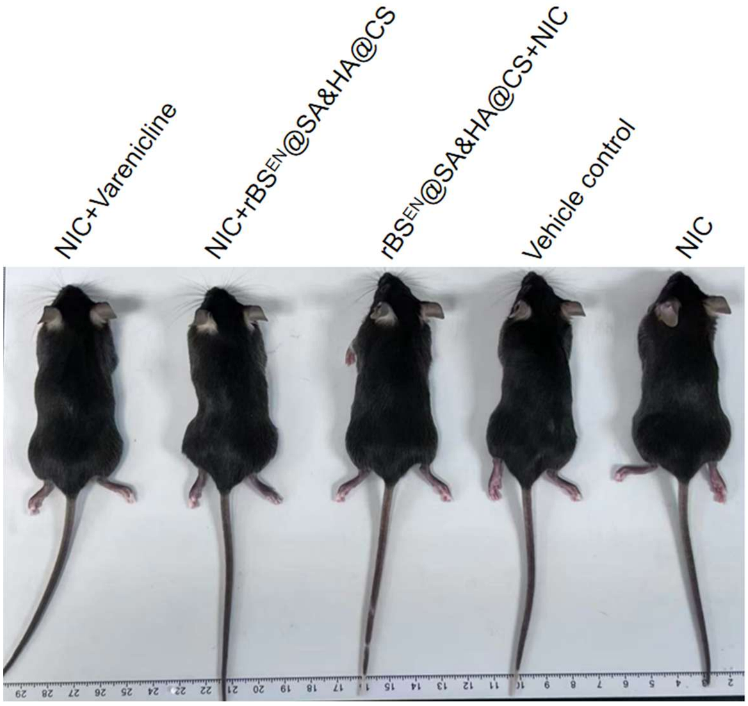


**Figure S25.** Representative images showing whisker loss in mice continuously exposed to nicotine.

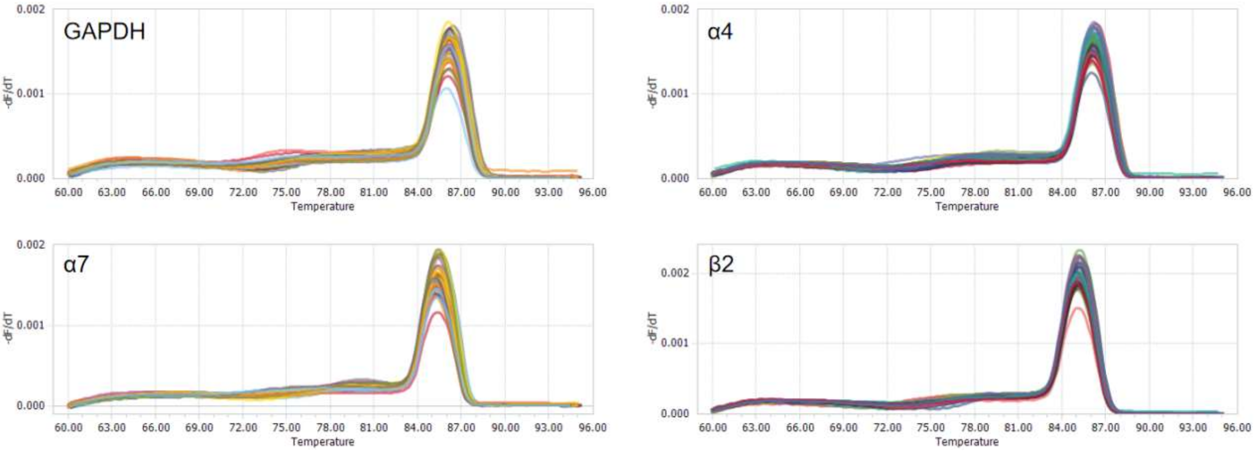


**Figure S26.** Melting curves of the primer pairs used for qPCR.

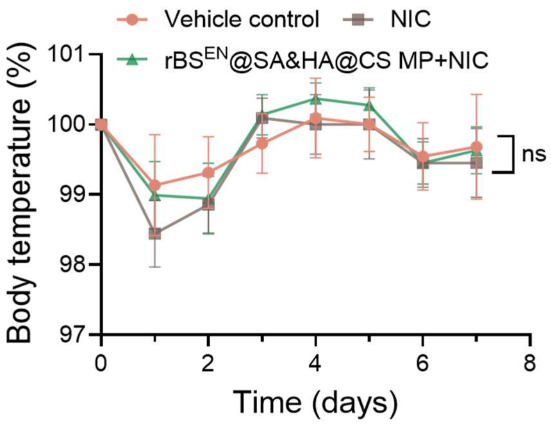


**Figure S27.** Body temperature changes among experimental groups in the nicotine toxicity assessments. Data are presented as mean ± SD (n = 6) and were analyzed using one-way ANOVA. ns, not significant.

#### Supporting Tables

**Table S1.** Summary of Keil's rules for trypsin miscleavage sites

| P4 | P3 | P2 | P1 | P1' | P2' | P3' |
|---|---|---|---|---|---|---|
| | | | [RK] | [P] | | |
| | | [WYF] | [RK] | [^RK] | | |
| | | [^RK] | [RK] | [WYF] | | |
| | | [DE] | [RK] | [^RK] | | |
| | | [^RK] | [RK] | [RK] | | |
| | | [^RK] | [RK] | [DE] | | |
| | | | [RK] | [^RK] | [DE] | [DE] |
| | | [^RK] | [RK] | [H] | | |
| [DE] | [DE] | [^RK] | [RK] | | | |
| | [DE] | [^RK] | [RK] | [^RK] | [DE] | |

*Different miscleavage patterns are expressed in the form of regular expressions. A set of letters within square brackets is treated as a single symbol, equivalent to any one of the characters inside the brackets, while the caret (^) denotes exclusion.

**Table S2.** Strains and plasmids used in this work

| Strains and plasmids | Description | Source |
|---|---|---|
| **Strains** | | |
| *E. coli* DH5α | Host strain for plasmid propagation and maintenance, *recA1* and *endA1* mutations | Sangon Biotech |
| *E. coli* BL21 (DE3) | Host strain for recombinant protein, T7 RNA polymerase gene | |
| *Bacillus subtilis* WB800N | *nprE aprE epr bpr mpr*::*ble nprB*::*bsr Δvpr wprA*::*hyg cm*::*neo; NeoR* | Bioscibio |
| *Bacillus subtilis* 168 | *trpC2* | |
| *B. subtilis* 168 S1 | Derived from *B. subtilis 168, ΔamyE*::*CotE-ssNox* | This work |
| *B. subtilis* 168 S2 | Derived from *B. subtilis 168, ΔgerKABC*::*CotE-ssNox* | |
| *B. subtilis* 168 S3 | Derived from *B. subtilis 168, ΔgerKABC*::*CotE-ssNox-dif* | |
| *B. subtilis* 168 S4 | Derived from *B. subtilis 168, ΔgerKABC*::*p43-CotE-ssNox-dif* | |
| *B. subtilis* 168 S501 | Derived from *B. subtilis 168, ΔgerKABC*::*CotE-ssNox-*$L_6$*-MnCAT-dif* | |
| *B. subtilis* 168 S502 | Derived from *B. subtilis 168, ΔgerKABC*::*CotE-ssNox-*$L_{15}$*-MnCAT-dif* | |
| *B. subtilis* 168 S503 | Derived from *B. subtilis 168, ΔgerKABC*::*CotE-MnCAT-*$L_{15}$*-ssNox-dif* | |
| *B. subtilis* 168 S6 | Derived from *B. subtilis 168, S502, ΔgerKABC::CotE-ssNox-*$L_{15}$*-MnCAT-CotY-ssNox-dif* | |
| **Plasmids** | | |
| pET32a (+) | *E. coli* expression vector, T7 promoter, *TrxA*, $Amp^r$ | Miaolingbio |
| pDG364 | *B. subtilis* integration vector, amyE arm, $Amp^r$, $Cm^r$ | Fenghbio |
| pDG364-01 | Derived from pDG364, *CotE-ssNox* inserted | This work |
| pDG364-02 | Derived from pDG364-01, amyE arm replayed with gerKABC | |
| pDG364-03 | Derived from pDG364-02, $Cm^r$ deletion Xer-cise plasmid | |
| pDG364-04 | Derived from pDG364-03, *promoter repalyed with P43* | |
| pDG364-0501 | Derived from pDG364-03, $L_6$*-MnCAT* inserted | |
| pDG364-0502 | Derived from pDG364-03, $L_{15}$*-MnCAT* inserted | |
| pDG364-0503 | Derived from pDG364-0502, *redesigned ssNox and MnCAT sites* | |
| pDG364-06 | Derived from pDG364-0502, *CotY-ssNox* inserted | |

**Table S3.** LC/MS/MS parameters for quantitative analysis by electrospray at positive mode

| | Q1 mass (Da) | Q3 mass (Da) | Q1 prerod offset (V) | CE (V) | Q3 prerod offset (V) |
|---|---|---|---|---|---|
| **Analytes** | | | | | |
| NIC | 163.15 | 130.15 | -18.0 | -19.0 | -25.0 |
| NMM | 161.10 | 119.10 | -18.0 | -27.0 | -21.0 |
| PON | 179.01 | 106.15 | -20.0 | -21.0 | -19.0 |
| **Internal standard** | | | | | |
| NIC-*d3* | 166.01 | 130.15 | -18.0 | -20.0 | -23.0 |

#### Supporting Sequences

***ssNox:***

ATGGATGAAAAAAGGAATAACGGACTATCACGCCGTTCGTTCATTGGCGGTGCGGCGG
TGGTGACCGCGGGTGCTGCTGGTTTGGGCCTTATCGGTAGCGCGAATGCTACTGAGAA
CGGCACCAGCAAGCGCGCGACCGGTTTCGACTACGACGTGATTGTTGTGGGTGGCGG
CTTCGCGGGCGCTACCGCGGCACGTGAGTGCGGTCATCAGGGTTATAAAACCCTGCTG
TTAGAAGCTAGATCTCGTCTGGGTGGTAGAACATTTACCAGCCATTTTGCCGGCCAAGA
AATTGAGTTCGGTGGAGCGTGGGTACATTGGCTGCAACCGCACGTGTGGAGCGAAATG
CAGCGCTACGGCTTGGGCGTGGTGGAGGACCCGCTGACGAACCTGGACAAGACGCTG
GTTATGTATAACGATGGATCTGTGGAGGACCTGCCGCCTGAAGTATTCGGCACGAATAT
CCAAGTCGCATTTGAAAAAATGTGCCACGACGCGTGGGAAGCATTTCCGAGGCCACA
CGAACCGATGTTTACCGAACGTGCACGTAAGCTGGACAAGATGAGCGTCCTGGACCGC
ATCAACCAGTTGGAGCTGACCCGTGCACAGCGTGCGGAGCTGAACTCCTATATGGCAT
TGTACGGTGGCGAAACAACCGACAAGTACGGGCTGCCGGGAGTGCTGAAATTGTTTG
CGTGTGGTGGTTGGAATTATAACGCGTTCATGGATACCGAAACCCACTACCGCATCGAA
GGCGGCACCATCGGCCTTATCAATGCTATGCTGGCAGATAGCGGCGCCGAGGTTCGCCT
CAACATGCCGGTTATTAGCGTGGAGCAGCTCAACGGTGGCGTTCGTGTGGAGACTGAC
GATGGTGAAACTATCACGGCCGGTACGATCATCATGACCGTTCCGCTGAACACCTATCG
CCATATTAACTTTACCCCGGCATTGTCCGAAGGCAAACAGCGTTTTATCCAAGAGGGTC
AGTTGTCGAAGGGTGCGAAACTCTACGTTCACGTTAAAGAAAACCTGGGCCGTGTTTT
CGCCTTTGCCGACGAACAACAACCGCTGAACTGGGTTCAGACCCACGATTATGGTGAT
GAACTGGGTACCATCTTATCCATTACCATCGCACGTGCGGAGACGATCGATGTAAATGA
TCGTGATGCGGTTACCCGTGAGATTCGTAAGTTATTCCCGGGTGTGGAGGTTTTGGGTA
TTGCCGCGTACGACTGGACCGCCGACCCGTTCAGCCTGGGCGCTTGGGCAGCTTACGG
TGTTGGTCAACTGAGTCGTCTGACCGACCTGCAGCAGCCAGAGGGCCGCATTTTGTTC
GCGGGCGCGGAAACTTCAAATGGTTGGCATGCTAATATTGATGGCGCGGTCGAGAGCG
GTCTGCGTGCGGGTCGCGAGGCGAAAGAGATTTTGGGCTAA

***P43* promoter:**

AGCTTCGTGCATGCAGGCCGGGGCATATGGGAAACAGCGCGGACGGAGCGGAATTTC
CAATTTCATGCCGCAGCCGCCTGCGCTGTTCTCATTTGCGGCTTCCTTGTAGAGCTCAG
CATTATTGAGTGGATGATTATATTCCTTTTGATAGGTGGTATGTTTTCGCTTGAACTTTTA
AATACAGCCATTGAACATACGGTTGATTTAATAACTGACAAACATCACCCTCTTGCTAA
AGCGGCCAAGGACGCTGCCGCCGGGGCTGTTTGCGTTTTTGCCGTGATTTCGTGTATC
ATTGGTTTACTTATTTTTTTGCCAAAGCTGTAATGGCTGAAAATTCTTACATTTATTTTAC
ATTTTTAGAAATGGGCGTGAAAAAAAGCGCGCGATTATGTAAAATATAAAGTGATAGCG
GTACCATTATAGGTAAGAGAGGAATGTACACATGAACAGACAAGAATTAATAACAGAA
GCT

***MnCAT:***

ATGTGGATTTATGAAAAAAAATTGCAATATCCTGTTCGTGTCAGCACATGCAACCCAAA
ATTGGCGAAATATTTAATTGAACAGTATGGAGGTGCGGATGGAGAACTCGCCGCTGCG
CTTCGTTATTTAAATCAACGCTACTCCATCCCTGATAAAGTAGTCGGTTTGCTCACAGAT
ATTGGAACCGAAGAATTAGCTCATTTGGAAATGATTGCGACAATGGTATACAAATTAAC
AAAAGACGCCACCCTAGAACAGCTGAAAGAAGCAGGTCTTGACGCACACTATGTCGA
TCATGACCGCGCTTTATTTTATCATAATGCCGCTGGAGTTCCATTTACGGCGACATACAT
TCAAGCGAAAGGAGACCCAATTGCCGATTTATATGAAGATATCGCTGCCGAAGAAAAA
GCGCGGGCAACCTATCAATGGATTATTAATATGAGCGATGATCCAGATTTAAACGATGG
GCTTCGCTTTTTACGTGAACGTGAAATCATTCATGCCCAGCGTTTTCGCGAAGCAGTCG
AAATTTTAAAAGAAGAGCGTGACCGCAAGAAAATTTTTTAG

***gerKABC:***

ATGCCTTTATTTTCTAAGCGAAAGAACAATACCGATTCAAAAGACAAGCAGAATACAG
ACGAAAGGAATCAGGAACAGCAGCAAGAAAAAGAAAGGCCGGTTCTCATTTCTCCAA
GTTTAGCAAAGAATATAGCGGAAACGAAAAAGGAAGTCGGAAGCAGCTCTGACGTCA
TCATTCGCGAGATTAAAATCGGGGAGCAGGATCATGTCCACCTAGCTGTTATCTATATTT
CTGGGCTGGTCGATAATAACACGATCCATGAGTCGTTAATTGACCCATTGGTACAGGAT
GAGTCTATCCAGAATACTCATGCGATCCAGCAAATTCTTGAAAAAAACGCTCCCGCTGG
GCGGGGTGAAAGCGGAAAAGAGCTGGGACAAGCTTTTTTCCGAGTTAATGCTTGGCA
ATGCGCTCATTTTTGCTGATGGCCATGATGAAGCTTTGATTTGCAGCACTCAAGGGGGA
GAGCAGCGCTCCATCCAAGAGCCGAGCACCCAGGTGTCATTCCGCGGCCCGCGCCAA
GGGTTTACAGAGTCTCTGCAAACCAATATTTCCATGATCCGGCGATACATTAAAAATCC
GAATTTATGGGTTGAAAAAATGAAAAAAGGTTCTGTTACGAATACGGATATCGCTCTCA
TGTACATTCAAGGAATATGTGATGAAAAGGTGCTGAAGGAAGTGAAACAGCGCCTGG
AAAAAATAGATATTGACAGTATTTTGGAATCTGGATATATTGAGCAGCTGATTGAAGATG
AAACCTTTACGACGTTTCCGACTATGTATCATACAGAACGCCCTGATGTTGTAGCGGGT
AATCTGTTAGAAGGAAGATTCGCGATTATTGTGGACGGAACACCGTTTGTGCTTATTGC
CCCAGCCTTATTTGTCCAGTTTTTTCAGTCTGTTGAGGATTATTATTCGCGTTTTGACATT
GCGACAAGCATAAGAATTCTTCGTGTTTTAGTCTTTTTCATTTCGCTTGTGGCACCAGC
TGTATACGTGGCAGCCACGACGTTTCATCAAGAAATGATACCGACTCAGCTGTTAGTGG
TCATTGCGGCCCAGCGGGAAATTGTCCCGTTTCCGGCTGTTGTGGAGGCGCTGACGAT
GGAGGTTGCTTTTGAGATTCTGAGGGAAGCGGGAGTCAGGCTGCCCCGGGTTGTCGG
CTCGGCAGTGTCAATCGTAGGCGCGCTCGTAATCGGCCAGGCTGCCGTACAGGCTGGT
ATCGTCTCTCCGGCAATGGTGATTATCGTGGCACTCACCGCGATTGCGAGCTTCGCAAC
ACCGGCGTTTGCAATGGCGATTTCAGCCCGGCTTATTCGGTTTATCTTTATTATTGCCTC
AGCTGTTATGGGTTTTTACGGGCTGATTTTAGGGATCATTATGATGTTTGTCCACTTATGC
AGTCTTCGTTCATTCGGCGTTCCATATATGTCGCCGCTTGCACCTTTTTCTTCACAAGGC
GTAAAGGACGCGCTTTTCCGGGTTCCGTGGTGGGCCGATGAAAAAAGGCCGGAATCG

GTCAGCAAGGAAGATAAGGTGCGGCAGGGGAAAGATCAACGCCCAGAGCCGGCCGCT
TCACGAGGAATGGTGAACAAAGATTTGGAAGAAGGGGATCAGAATGGTACGTAAATG
TCTATTAGCCGTTCTCATGCTTTTGAGCGTCATTGTGCTGCCCGGCTGCTGGGATAAGC
GGGAGCTGACGGATCTTGCGATTATCTCGGCGATTGGCATCGATCGGACAAACGACAG
CAACTACGTGCTGCATCTTCAAATTATCAATCCAGGAAATGTTGCCGGCGGCCTTCAGG
GAGGCGGTGCTGGAGACAGGCCGCCTGTATCTGTCTATTCGATTGAAGGTAACAATATA
ACGGAAGCACTCAGAAAAGCCTCCATGAAGGTATCACGGCGGCTTTATTTTGCCCATA
CCAATCTAGTCGTGATCAATGAAAAGCTGGCAAAAGAAGAAGGTTTGGATTTTGTATTA
GATAATCTTGACCGGGACACAGAATTCAGGACAACAGCAACATTCGTCGTTGCCCATA
AAACAAAAGCAGAAAATATTGTGAAAATCTTGACGCCGATTGATAAAATTCCGTCAAA
TAAAGTCAATAAAACACTCGATTTCACGGAGGCACAATACGGGCGGGTGGTCAAAATC
AATATTCAGGACGTTTTGAAAACACTTGCCGCCAACACTATGGCACCGGTCATCCCCG
GATATATGATGATTGGTGACGATAAAAAAGGAGTCAGCATGGAAAACACGCAGGCGAC
TGACCCGAAAGCCATTCTTCAGGCAGATGGTTTGGCGGTTTTTGATAAGGCAGGATATT
TAAAGTATTGGCTGGAAGATGATGAAAGCGTGGGGGCAGTCTGGCTTATGAATAAGAT
CCAGCATACGTTTATTAATGCTGACTGGGGCAAGACGAAGGATGCGGTCAGCTTGCAG
GTTACCCATCAAGATACAAAGCTGGTACCGAAAATGCGGAACGGACGCCCGTACATCC
ATGTGAAAGTGTCGGTTGAAGGTATTATAGATGCTGTCAAATACCCTTTCCAATTGTCTG
ATCCCAAGGTGCTGGCCGCTATTGAAAAAGCGCTCAACAAAGAGCTTGAAAAGGAAA
TCAGCCATACTGTGAAAAAGATTAAGAAGAATAAAATAGATTTTATCGGTTTTGGTGAC
ACGATATATAGAAAGTATCCGGAGCAGTGGGAGAAAATGAAGGACACTTGGGACAAA
GAATATTTGCCGGAACTGCCCATCGATGTGAAGGCCGAGACGTATATCAGAAGGACGG
GATTGCGAAACAATCCGATCAAACACCAGTTTAAAGATGATTAGGTCTGAAGAAAGGG
GAAGTGACAGATGGAAAAAGCCAGAATAAGCATAAGGCAGTTGTTTGTCATGATTATC
ATTTTTGAACTGGGCAGCTCCTTATTGATTACACCGGGATCAATGGCGGGCAGGGATGC
TTGGATAGCAGTTTTATTAGGCTGTGCGATCGGACTGTTTCTTTTCTATTTGTATCAAGG
CATTTATCAATGTTATCCGAATTCTTCTCCGAAAGAATATATGGATGATATGCTGGGAAC
CAAGCTGAGCTGGCTGTTTTCATTTCTCTATATCTTGTACTTTGCCTATATTGCCGCAAG
GGTGCTGCGTGACTTTGGAGAAATGCTGCTGACGTTTGCTTATCATGATACACCTATCAT
TATTGTAAATGCTTTATTAATGGTAGTAAGCATATATGCCGTAAGAAAAGGAATTGAAGT
ACTCGCACGCGCGGCCGAACTTCTTTTCGGGGCCATGTATTTACTGGGCGCGATCGGCC
TTGTGCTGATTATTGTCTCAGGGACTATAGACCCGCACAATTTAAAGCCGGTTTTGGCA
AACGGCATTTCTCCTGTCCTTCACTCTGTTTTCACACAGACCATGTACGTTCCATTTGGT
GAAGTTGTTTTATTTGTGATGATTTTCCCTAACCTGAATGACAGAAAAGATGTGAAAAA
AATGGGGATGATCGCCATGGCTATTAGCGGATTGATTGTGGCGCTCACCGTGGCGATTA
ATATTAGCGTGCTTGATGTTGATCTTACACTCAGATCCCAGTTTCCGCTTTTAAGCACGA
TCCAGACAATCAAGGTCGAAGAATTTTTAGACCGGCTCGATGTCTTTTTTATGCTGGCG
CTGATCATCGGCGGCTTCTTTAAGGTCAGCCTCTACTTATATGCCACTGTGGTTGGGAC

ATCTACGCTCTTTAAGGAAAAGAACCCCTCTCAATTGGCTTATCCGATGGGATTGGGCA
TCTTAATCCTTTCTATAACGATCGCGACTAACTTTTCGGAGCACTTGAATGAAGGCCTG
AACGTGGTGCCGCTGTATATTCATTTGCCTTTTCAATTATTGTTTCCGCTCTTTCTGTTTA
TTGTGGCCGTTTGGAAAAAGAAACGAAGAGAGAAGTCAAAAGGGGAGGAAGCAAAA
AAATGA

***amyE:***

ATGTTTGCAAAACGATTCAAAACCTCTTTACTGCCGTTATTCGCTGGATTTTTATTGCTG
TTTCATTTGGTTCTGGCAGGACCGGCGGCTGCGAGTGCTGAAACGGCGAACAAATCGA
ATGAGCTTACAGCACCGTCGATCAAAAGCGGAACCATTCTTCATGCATGGAATTGGTCG
TTCAATACGTTAAAACACAATATGAAGGATATTCATGATGCAGGATATACAGCCATTCAG
ACATCTCCGATTAACCAAGTAAAGGAAGGGAATCAAGGAGATAAAAGCATGTCGAACT
GGTACTGGCTGTATCAGCCGACATCGTATCAAATTGGCAACCGTTACTTAGGTACTGAA
CAAGAATTTAAAGAAATGTGTGCAGCCGCTGAAGAATATGGCATAAAGGTCATTGTTG
ACGCGGTCATCAATCATACCACCAGTGATTATGCCGCGATTTCCAATGAGGTTAAGAGT
ATTCCAAACTGGACACATGGAAACACACAAATTAAAAACTGGTCTGATCGATGGGATG
TCACGCAGAATTCATTGCTCGGGCTGTATGACTGGAATACACAAAATACACAAGTACA
GTCCTATCTGAAACGGTTCTTAGACAGGGCATTGAATGACGGGGCAGACGGTTTTCGA
TTTGATGCCGCCAAACATATAGAGCTTCCAGATGATGGCAGTTACGGCAGTCAATTTTG
GCCGAATATCACAAATACATCTGCAGAGTTCCAATACGGAGAAATCCTGCAGGATAGTG
CCTCCAGAGATGCTGCATATGCGAATTATATGGATGTGACAGCGTCTAACTATGGGCATT
CCATAAGGTCCGCTTTAAAGAATCGTAATCTGGGCGTGTCGAATATCTCCCACTATGCAT
CTGATGTGTCTGCGGACAAGCTAGTGACATGGGTAGAGTCGCATGATACGTATGCCAAT
GATGATGAAGAGTCGACATGGATGAGCGATGATGATATCCGTTTAGGCTGGGCGGTGAT
AGCTTCTCGTTCAGGCAGTACGCCTCTTTTCTTTTCCAGACCTGAGGGAGGCGGAAAT
GGTGTGAGGTTCCCGGGGAAAAGCCAAATAGGCGATCGCGGGAGTGCTTTATTTGAAG
ATCAGGCTATCACTGCGGTCAATAGATTTCACAATGTGATGGCTGGACAGCCTGAGGA
ACTCTCGAACCCGAATGGAAACAACCAGATATTTATGAATCAGCGCGGCTCACATGGC
GTTGTGCTGGCAAATGCAGGTTCATCCTCTGTCTCTATCAATACGGCAACAAAATTGCC
TGATGGCAGGTATGACAATAAAGCTGGAGCGGGTTCATTTCAAGTGAACGATGGTAAA
CTGACAGGCACGATCAATGCCAGGTCTGTAGCTGTGCTTTATCCTGATGATATTGCAAA
AGCGCCTCATGTTTTCCTTGAGAATTACAAAACAGGTGTAACACATTCTTTCAATGATC
AACTGACGATTACCTTGCGTGCAGATGCGAATACAACAAAAGCCGTTTATCAAATCAAT
AATGGACCAGAGACGGCGTTTAAGGATGGAGATCAATTCACAATCGGAAAAGGAGATC
CATTTGGCAAAACATACACCATCATGTTAAAAGGAACGAACAGTGATGGTGTAACGAG
GACCGAGAAATACAGTTTTGTTAAAAGAGATCCAGCGTCGGCCAAAACCATCGGCTAT
CAAAATCCGAATCATTGGAGCCAGGTAAATGCTTATATCTATAAACATGATGGGAGCCG
AGTAATTGAATTGACCGGATCTTGGCCTGGAAAACCAATGACTAAAAATGCAGACGGA
ATTTACACGCTGACGCTGCCTGCGGACACGGATACAACCAACGCAAAAGTGATTTTTA

ATAATGGCAGCGCCCAAGTGCCCGGTCAGAATCAGCCTGGCTTTGATTACGTGCTAAAT
GGTTTATATAATGACTCGGGCTTAAGCGGTTCTCTTCCCCATTGA

***CotB* with promoter:**

ACGGATTAGGCCGTTTGTCCTCATGGACCCGTATAAAAAGAATGATATTGAGCGTTTTG
ACCGTGAGCCGGATGTGATCTGCGAGTATATTAAAAACCGTTCACAATACCTCAAAGAT
CATTTAAGTATTTTATGAATGCGTGAAAATGGGTATTCGCGGAAAAAGCGACAATTAGG
CTATTGAATTAGTTCAACAAATAAATGTGACACGTATATATGCAGTATGTTTATCATCTAT
GTATAAGTGACTAGGAGGAATTTGAATGAGCAAGAGGAGAATGAAATATCATTCAAATA
ATGAAATATCGTATTATAACTTTTTGCACTCAATGAAAGATAAAATTGTTACTGTATATCG
TGGAGGTCCGGAATCTAAAAAAGGAAAATTAACAGCTGTAAAATCAGATTATATAGCTT
TACAAGCTGAAAAAAAAATAATTTATTATCAGTTGGAGCATGTGAAAAGTATTACTGAG
GATACCAATAATAGCACCACAACAATTGAGACTGAGGAAATGCTCGATGCTGATGATTT
TCATAGCTTAATCGGACATTTAATAAACCAATCAGTTCAATTTAACCAAGGGGGTCCGG
AATCTAAAAAAGGAAGATTGGTCTGGCTGGGAGATGATTACGCTGCGTTAAACACAAA
TGAGGATGGGGTAGTGTATTTTAATATCCATCACATCAAAAGTATAAGTAAACACGAGC
CTGATTTGAAAATAGAAGAGCAGACGCCAGTTGGAGTTTTGGAAGCTGATGATTTAAG
CGAGGTTTTTAAGAGTCTGACTCATAAATGGGTTTCAATTAATCGTGGAGGTCCGGAAG
CCATTGAGGGTATCCTTGTAGATAATGCCGACGGCCATTATACTATAGTGAAAAATCAAG
AGGTGCTTCGCATCTATCCTTTTCACATAAAAAGCATCAGCTTAGGTCCAAAAGGGTCG
TACAAAAAAGAGGATCAAAAAAATGAACAAAACCAGGAAGACAATAATGATAAGGAC
AGCAATTCGTTCATTTCTTCAAAATCATATAGCTCATCAAAATCATCTAAACGATCACTA
AAATCTTCAGATGATCAATCATCC

***CotC* with promoter:**

GGATAAATCGTTTGGGCCGATGAAAAATCGGCTCTTTATTTTGATTTGTTTTTGTGTCAT
CTGTCTTTTTCTATCATTTGGACAGCCCTTTTTTCCTTCTATGATTTTAACTGTCCAAGCC
GCAAAATCTACTCGCCGTATAATAAAGCGTAGTAAAAATAAAGGAGGAGTATATATGGG
TTATTACAAAAAATACAAAGAAGAGTATTATACGGTCAAAAAAACGTATTATAAGAAGT
ATTACGAATATGATAAAAAAGATTATGACTGTGATTACGACAAAAAATATGATGACTATG
ATAAAAAATATTATGATCACGATAAAAAAGACTATGATTATGTTGTAGAGTATAAAAAGC
ATAAAAAACACTAC

***CotE* with promoter:**

CCGGTTCAAAGGTGAAGCATTCAAATAACAGCTGTTCTCTCTAAACACGGTGCCTTTAC
AGGCCCGTGTTTTTTTATCATTTGTGCGGTTAAAAATGAACTAAATAATCTATGTACCAA
ATGTTCAATTGGTTTTTCTGTGCTCAGCCGCGTATAAACTTTATCGCACTTATAAGTAAA
GTTTCTAGGCACCCCTGCATACAATGGAACAGAAACTTTGTATTTTTATATTTTATTTATA
AAAATGCACACTAGACAAATGCCCAGCATAAGATAACACGAAGAAGAACAAGGAGGC
ATGCCGGAATGTCTGAATACAGGGAAATTATTACGAAGGCAGTAGTAGCGAAAGGCCG
AAAATTCACCCAATGCACCAACACCATCTCGCCTGAGAAAAAACCGAGCAGCATTTG

GGTGGTTGGATTATTAACCACAAGTATGACGCTGAAAAAATTGGAAAAACGGTAGAAA
TTGAAGGGTATTATGATATAAACGTATGGTACTCTTACGCGGACAACACAAAGACAGAG
GTTGTCACAGAACGGGTAAAATATGTAGATGTCATTAAACTCAGATACAGAGACAATAA
TTACTTAGATGATGAGCATGAAGTGATTGCCAAAGTGCTTCAGCAGCCAAACTGCCTT
GAAGTGACCATTTCGCCGAATGGAAATAAAATCGTTGTGCAGGCAGAAAGAGAATTTT
TGGCGGAAGTGGTAGGGGAAACAAAGGTAGTTGTTGAGGTCAATCCTGACTGGGAAG
AGGATGACGAGGAAGATTGGGAAGATGAGCTTGATGAAGAGCTTGAAGACATCAACC
CGGAGTTTTTAGTGGGAGATCCTGAAGAA

***OxdD* with promoter:**

CTGAAGATGAAGTGATAGGTCATGATGTATTAGTAAAAGTGGAGTGGAGGACAGGCCA
GAAAAAACAGACAGAAGCAATCAAATTACATAAGAAGCCATGGTATAAAAAATAGTTT
ATTTGATGTATTTGTGATCACATTGGTGGTCACTTTTTTATTTGCGGATTCCTAGGCACA
GCAATCTAAGATTCTGCATAGGCTGAAATAAAATCTTGTTCATTTCTAAAACGAGGTGC
ATGCTGTTGGAACAACAACCAATCAATCATGAAGACAGAAACGTGCCGCAGCCTATTC
GAAGTGATGGAGCTGGAGCTATTGATACAGGCCCGCGAAATATAATACGGGATATTCAA
AATCCGAATATATTTGTTCCGCCTGTTACAGATGAGGGTATGATTCCTAACTTGAGATTT
TCATTCTCAGACGCTCCCATGAAATTAGATCACGGCGGCTGGTCAAGAGAAATCACCG
TAAGACAGCTTCCGATTTCGACTGCGATTGCAGGTGTAAACATGAGCTTAACTGCGGG
AGGCGTCCGCGAGCTTCATTGGCATAAGCAAGCGGAGTGGGCTTATATGCTTTTGGGA
CGGGCACGTATCACCGCTGTTGACCAAGACGGACGAAATTTCATTGCTGATGTTGGTC
CCGGCGACCTTTGGTACTTCCCGGCAGGAATTCCGCATTCCATACAGGGATTGGAACA
CTGCGAGTTTCTGCTCGTTTTCGATGATGGGAACTTTTCTGAGTTTTCAACGTTAACCA
TTTCAGATTGGCTTGCACACACACCAAAAGATGTTCTGTCTGCAAATTTCGGTGTCCCG
GAGAATGCTTTCAACTCTCTTCCGTCTGAGCAAGTCTATATCTACCAAGGGAATGTGCC
GGGATCAGTCGCCAGTGAAGACATTCAGTCACCATATGGAAAAGTCCCCATGACCTTT
AAACACGAGCTGTTAAATCAACCCCCAATTCAAATGCCAGGGGGAGTGTACGAATTGT
GGATTCTTCTAACTTCCCAATTTCAAAAACGATAGCCGCTGCACTTGTTCAGATTGAGC
CTGGCGCGATGAGAGAGCTTCATTGGCATCCCAATAGCGATGAGTGGCAATATTATCTA
ACAGGACAGGGACGAATGACGGTATTTATCGGAAATGGGACTGCCCGCACATTTGATT
ATAGAGCAGGCGACGTTGGATACGTGCCTTCTAATGCCGGACACTATATACAAAACACT
GGTACAGAAACATTATGGTTTTTAGAAATGTTCAAAAGTAACCGCTATGCAGATGTGTC
ACTCAATCAGTGGATGGCATTGACGCCTAAAGAATTAGTACAAAGCAACTTGAATGCT
GGATCAGTCATGCTTGATTCTCTGCGCAAGAAGAAAGTGCCTGTTGTGAAATATCCCGG
TACG

***CotG* with promoter:**

TTTGTATTTCTTTTTGACTACCCAGCAATTGCCGTCTTTCCAGTATTCTTTTTTGCTGTCA
TAATCATCGTGTCTTTTGTAGTCGTCGCAGTGGTGGTGCGGTTTTTTGTAGTATTTCTTAT
GGGAACGATAAGATTTTTGTGTGTGCGGCATGACTTCTTATAGCTGCGAGATTTTTTCT

TGTGAGAACAGTAAGACTTTTTGTAGCTGCGAGATTTTTTGTAAGAACGGCAAGATTTT
TTATAGCTACGAGATTTTTTGTAAGAACGGTAAGATTTTTTATAGCTGCGAGATTTTTTG
TGAGAACGGTACGATTTTTTGTGGCTGCGAGATTTCTTGTGAGAGCAGTATGATTTTTT
GTGGCTGCGAGATTTTTTGTGAGAACAGAATGATTTTTTGTGACTGCGAGATTTTTTGT
GAGAGCAGTATGATTTTTTATGGCTGCGAGATTTTTTGTGAGTGCGGTGCGACTTTTTAT
GACTGCGTTTTTTTCCATGAGGCTCTTGGTATAAATAATCCTTTAAACCTTCTTTTTTTGC
GGATTTCACCGCTTCTTCGATGTCAGAATGGGAATAGTGGCCCAATTTGAAATCCTCCT
TTTAAAAATATGTTAAGGATGAAGATCCTTACTACCCTCATAGTACGCGCGGTTTTAAAA
AGTGTATAAGTGTTCCGCTGAGATCTTATAAAAATAAAAGCCTAAAATTTGTAAGGAGG
ATTATGGATGAAGAATCAATCCAATTTACCGCTTTATCAGCTGTTTGTTCATCCAAAAGA
CTTGCGTGAATTAAAAAAGGATATATGGGACGATGATCCGGTGCCAGCTGTGATGAAG
GTAAATCAAAAAAGGCTGGATATTGATATCGCTTATCGGGGATCACATATCAGAGACTT
CAAAAAGAAGTCATACCATATTTCCTTTTATCAGCCGAAAACATTCCGCGGCGCGCGAG
AGATTCACTTAAATGCGGAGTATAAAGACCCTTCCTTGATGAGAAACAAATTGTCTCTG
GATTTTTTCTCGGAGCTAGGGACAC

***CotX* with promoter:**

CCAATGACGAAAATGCTGGTCCAAAATATATTTTCTAAACATGGGGTCACAAAAGACA
AAATGAAAAAAGTATCAGACGAAGAAAAAGAAATGCTCTTAAATCTTGTAAAAGACTT
ACAAGCTAAATCACAAGCGCTAATAGAAAACCAAAAGAAGAAAAAAGAAGAAGCAG
CCGCACAAGAGCAAAAGAACACAAAACCGTTAAGCCGCAGAGAGCAGCTCATTGAAC
AGATCAGACAAAGACGGAAAAACGATAACAATTAGCCAATCATAAAAAATAGGGTTCT
TCATCAGGATATATGACTCAGTCAAAATAAGAGGCTCGCTCATTTAATAACAGTAAAAG
AAAAGGAGGAATAGATATGGAAAGCAGACCATATTCTTGGGTTGCACTTGATCCTGACT
GTGACCATCCGTTAGATGACAAAGAGAAAGATAAAGAAAAACACGAAAGAAAATGTC
ATTGCGACGTTTGCTGTAATGGCAATGGTTTTTTTGGCAACGACAACGCCTTCATCGAC
CAAGATCTAGCTCAAGCAAATCTCAACAAACAAGTTTCAGATGAAACGATTATTATTAG
AGATTCTTGTGACATCAATGTTACATCTACAGACGTTCAAGCCGTAACATCAGTTGTAA
CAGCACTTAATGCCGCTGTCGTAACGGCAACTCTGACATCAATTGCAGACGGCGTAATT
GCCGAATTAGTCGCACAAGATTTGTTACAGCTTACAGCTAACAAACAAGTAAACCGCC
AAAAACTTCTCATCGAATGTTCCCGCGGCGTAAACGTCACAACAGTAGATGCCGATATC
GCAACCCTTATTTCTACAGCAACAAATACACTCGTAGCCATCCTAGTTATCACTCTTGTC
CTC

***CotY* with promoter:**

GCAGAGCTAAAAACGCTCTGCTTTTTCTTATTTTCCAAGCATATGATGAATATATAGACG
TTCACCCACACCAAGTGGGGCACGGGTACATATGTTGTTAAGGACTAAAGTCAAATAC
CCTATAAAGAAGGAGCTGAAATCAATGAGCTGCGGAAAAACCCATGGCCGGCATGAG
AACTGTGTATGCGATGCAGTGGAAAAGATTTTAGCAGAGCAGGAGGCAGTTGAAGAA
CAGTGTCCGACTGGCTGCTATACCAACCTTTTAAACCCTACGATTGCTGGAAAAGACA

CAATTCCGTTTCTCGTTTTTGATAAAAAAGGCGGATTGTTCTCCACATTCGGAAACGTA
GGGGGATTTGTGGATGATATGCAATGCTTTGAATCCATTTTCTTCCGCGTCGAAAAATTA
TGCGATTGCTGTGCAACACTGTCTATTTTACGCCCGGTCGATGTCAAAGGCGATACCTT
AAGTGTTTGCCACCCTTGCGACCCGGATTTCTTCGGGCTAGAAAAAACAGATTTCTGC
ATTGAAGTGGATCTCGGATGCTTCTGCGCGATTCAGTGCCTGTCACCAGAGCTAGTTGA
CAGAACATCGCCTCACAAAGATAAAAAGCATCATCACAATGGA

***CotZ* with promoter:**

CCTAGTTATCACTCTTGTCCTCTAGGACCTAAAAGCAGAGCTAAAAACGCTCTGCTTTT
TCTTATTTTCCAAGCATATGATGAATATATAGACGTTCACCCACACCAAGTGGGGCACG
GGTACATATGTTGTTAAGGACTAAAGTCAAATACCCTATAAAGAAGGAGCTGAAATCAA
TGAGCTGCGGAAAAACCCATGGCCGGCATGAGAACTGTGTATGCGATGCAGTGGAAA
AGATTTTAGCAGAGCAGGAGGCAGTTGAAGAACAGTGTCCGACTGGCTGCTATACCAA
CCTTTTAAACCCTACGATTGCTGGAAAAGACACAATTCCGTTTCTCGTTTTTGATAAAA
AAGGCGGATTGTTCTCCACATTCGGAAACGTAGGGGGATTTGTGGATGATATGCAATGC
TTTGAATCCATTTTCTTCCGCGTCGAAAAATTATGCGATTGCTGTGCAACACTGTCTATT
TTACGCCCGGTCGATGTCAAAGGCGATACCTTAAGTGTTTGCCACCCTTGCGACCCGG
ATTTCTTCGGGCTAGAAAAAACAGATTTCTGCATTGAAGTGGATCTCGGATGCTTCTGC
GCGATTCAGTGCCTGTCACCAGAGCTAGTTGACAGAACATCGCCTCACAAAGATAAAA
AGCATCATCACAATGGATAAACACTTGTAAAGAGGAATCCCCTTCCAAACTCTATTACA
GGTGCAAATCATTCAGCTGCTTCCAGCGTGAGGCAGCTGACTCTCCATCATTTCGCCAG
TCTCAAATTTATAAGGCAGGAGGGATAATATGAGCCAGAAAACATCAAGCTGCGTGCG
TGAAGCTGTAGAAAATATTGAAGATCTGCAAAACGCTGTTGAAGAAGACTGCCCGACC
GGCTGCCACTCTAAGCTTTTATCTGTAAGCCATTCGTTAGGCGACACAGTGCCTTTTGC
AATATTTACATCAAAATCAACGCCATTAGTCGCCTTCGGAAATGTCGGCGAACTCGATA
ACGGCCCTTGCTTTAATACAGTATTTTTCAGGGTCGAAAGAGTGCATGGAAGCTGTGCA
ACACTGTCATTATTAATCGCATTTGACGAACACAAACACATTTTGGACTTCACCGATAA
AGATACGGTGTGTGAAGTGTTCCGACTCGAAAAAACGAACTACTGTATTGAAGTTGAC
TTAGACTGCTTCTGCGCAATCAACTGCTTAAATCCTCGATTAATCAATCGTACACATCAT
CAT

#### SI References


1. Chen C *et al*. TBtools-II: A "one for all, all for one" bioinformatics platform for biological big-data mining. *Mol Plant* **16**, 1733–1742 (2023).
2. Siepen JA, Keevil EJ, Knight D, Hubbard SJ. Prediction of missed cleavage sites in tryptic peptides aids protein identification in proteomics. *J Proteome Res* **6**, 399–408 (2007).
3. Gershon PD. Cleaved and missed sites for trypsin, lys-C, and lys-N can be predicted with high confidence on the basis of sequence context. *J Proteome Res* **13**, 702–709 (2014).
4. Abramson J *et al*. Accurate structure prediction of biomolecular interactions with AlphaFold 3. *Nature* **630**, 493–500 (2024).
5. Niu C *et al*. Engineering of yersinia phytases to improve pepsin and trypsin resistance and thermostability and application potential in the food and feed industry. *J Agric Food Chem* **65**, 7337–7344 (2017).
6. Xiong X *et al*. Engineering *Bacillus* spores to display nicotine oxidase: *in situ* specific and sensitive nicotine detection. *ACS Sens* **10**, 3589–3599 (2025).
7. Nguyen HD, Phan TT, Schumann W. Analysis and application of *Bacillus subtili*s sortases to anchor recombinant proteins on the cell wall. *AMB Express* **1**, 22 (2011).
8. Hinc K *et al*. Expression of genes coding for GerA and GerK spore germination receptors is dependent on the protein phosphatase PrpE. *J Bacteriol* **188**, 4373–4383 (2006).
9. Bloor AE, Cranenburgh RM. An efficient method of selectable marker gene excision by Xer recombination for gene replacement in bacterial chromosomes. *Appl Environ Microbiol* **72**, 2520–2525 (2006).
10. Akinola LS *et al*. C57BL/6 substrain differences in pharmacological effects after acute and repeated nicotine administration. *Brain Sci* **9**, 244 (2019).
11. Kim B, Im HI. Behavioral characterization of early nicotine withdrawal in the mouse: a potential model of acute dependence. *Behav Brain Funct* **20**, 1 (2024).
12. Han Y *et al*. Cysteine sulfenylation contributes to liver fibrosis via the regulation of EphB2-mediated signaling. *Cell Death Dis* **15**, 602 (2024).
